# Loss of hand control expressiveness revealed by task- and individual-specificity in spatiotemporal finger coordination

**DOI:** 10.64898/2026.03.30.715145

**Authors:** Patrick Ihejirika, Divya Rai, Michael Rosenberg, Jing Xu

## Abstract

Stroke impairs dexterous hand use in daily activities, with prior studies of endpoint postures demonstrating reduced coordination complexity and diminished task-appropriate and individually-distinctive (termed *expressiveness*) coordination required by these daily activities. However, how spatiotemporal coordination reveals individual differences in post-stroke coordination remains unclear. Here, we characterized spatiotemporal coordination in able-bodied and post-stroke hands during finger individuation. We quantified coordination complexity and expressiveness (task-, individual-, and group-specificity) using principal component analysis (PCA) and linear discriminant analysis of 3D isometric forces from all five fingers. Paretic fingers showed reduced complexity and expressiveness, which was associated with greater intrusion of flexor bias in the paretic hand. Higher-variance PCs were characteristic of tasks and groups, while both higher- and lower-variance PCs were characteristic of individual-specific coordination. These findings suggest that stroke reduces spatiotemporal finger coordination complexity and expressiveness, limiting the coordination patterns available for daily hand use.

## Introduction

Despite well-documented impairments in finger coordination after stroke, the spatiotemporal organization of finger coordination remains poorly understood. Impaired coordination limits execution of activities of daily living that require precise, time-varying modulation of forces across multiple fingers, such as manipulating small objects or using hand tools; and it also compromises a person’s expression of their individuality. Compared to able-bodied adults, stroke survivors show reduced ability to selectively coordinate (i.e., individuate) fingers isotonically (<u>Lang & Schieber, 2003</u>, <u>2004</u>; <u>Schieber et al., 2009</u>) and isometrically (i.e., enslavement; <u>Li et al., 2003</u>; <u>Xu et al., 2017</u>), along with finger flexor bias intrusion (<u>Xu et al.,</u> <u>2017</u>). However, these studies quantified coordination using endpoint postures and may fail to capture how the fingers are coordinated over time while performing a task. Indeed, endpoint forces can be achieved using several coordination strategies, such that endpoint coordination may not fully reflect individual differences in post-stroke impairments (<u>Bernstein., 1967</u>; <u>Levin et</u> <u>al., 1996</u>; <u>Valero-Cuevas et al., 1998</u>; <u>Valero-Cuevas., 2000</u>; <u>Levin et al., 2002</u>; <u>van Kordelaar et</u> <u>al., 2013</u>). Characterizing the spatial and temporal aspects of finger coordination may further reveal individual differences in post-stroke coordination strategies underlying finger movement that are not apparent when analyzing only endpoints. Identifying maladaptive spatiotemporal finger coordination patterns may reveal novel, impairment-specific treatment targets.

Across naturalistic tasks, the human hand exhibits remarkable coordination complexity that can be revealed using dimensionality reduction techniques, but spatiotemporal coordination has yet to be explored. Early studies of constrained grasping suggested that most of the variance in hand postures could be explained by two or three principal components (PCs) (<u>Santello et al.,</u> <u>1998</u>; <u>Santello & Soechting, 2000</u>; <u>Jarque-Bou et al., 2019</u>). In contrast, later studies demonstrated that finger coordination complexity is not fixed but rather depends on task demands and behavioral context, with more naturalistic movements requiring additional PCs to explain comparable variance in hand postures (<u>Todorov & Ghahramani, 2004</u>; <u>Ingram et al.,</u> <u>2008</u>; <u>Thakur et al., 2008</u>). More recent studies show that *subtle components* of coordination (i.e., low-variance PCs) capture meaningful behavioral information: PCs accounting for 1% of the variance in hand postures can distinguish grasp types and American Sign Language gestures (<u>Yan et al., 2020</u>) and encode task-specific features like finger rotation direction of simulated kinematics (<u>Chiappa et al., 2024</u>). Aligning with these findings, we previously showed that post-stroke fingers exhibit reduced coordination complexity, requiring fewer PCs to explain comparable endpoint force variance and losing variance in finger-coactivation shape during individuated force production than able-bodied fingers (<u>Xu et al., 2023</u>). Collectively, these findings demonstrate that the complexity of finger coordination reflects task demands and neural constraints, and that low-variance PCs may encode subtle but task-relevant information. However, what these different aspects of finger coordination represent remains unclear.

Task-, individual-, and group-specific aspects of finger coordination may reveal how stroke disrupts coordination. In able-bodied fingers, coordination varies across tasks, with even subtle aspects of finger posture contributing to reliable distinctions between behaviors (<u>Yan et al.,</u> <u>2020</u>). Such *task-specific* information likely reflects the ability to flexibly combine control dimensions to produce distinct movements. Coordination is also *individual-specific*, likely shaped by lifelong motor experiences and learning (<u>Ting et al., 2024</u>) and evident across behaviors from gait (<u>Winner et al., 2023</u>) to culturally shaped hand movements, such as pottery (<u>Gandon et al., 2013</u>). We refer to the hand’s capacity to generate diverse, task-appropriate, and individually-distinctive coordination patterns as *expressiveness*. Stroke reduces the independence of muscle activations (<u>Dewald et al., 1995</u>), and exaggerates flexor patterns, potentially limiting access to subtle-but-important neural control strategies encoded in lower-variance PCs (<u>Xu et al., 2023</u>). Consequently, post-stroke changes to coordination may reduce finger flexibility, making coordination less distinct across tasks and individuals, and may also exhibit *group-specific* changes characteristic of paretic fingers. However, these changes likely vary across individuals due to differences in lesion characteristics and compensatory mechanisms (<u>Lodha et al., 2010</u>, <u>2013</u>). Determining whether post-stroke coordination remains distinguishable across tasks and individuals may provide insight into how stroke reshapes the expressive capacity of finger coordination.

Compared to finger kinematics, isometric fingertip forces may better capture individual differences in the neural control of finger coordination. While kinematic analyses describe finger motion, similar postures can be produced by different combinations of joint moments and muscle activation patterns, limiting inference about underlying neural and biomechanical constraints from kinematics alone (<u>Hollerbach & Flash, 1982</u>; <u>Hoffman & Strick, 1986</u>; <u>Zajac &</u> <u>Gordon, 1989</u>; <u>Valero-Cuevas et al., 1998</u>). In contrast, isometric forces may more directly reflect individual differences in these constraints, as muscle activations are more closely tied to fingertip forces than finger positions (<u>Nowak, 2008</u>; <u>Johansson & Westling, 1988</u>). Additionally, isometric force production tasks also mitigate posture-dependent biomechanical factors like muscle and tendon lengths on force production (<u>Zajac., 1989</u>). Prior studies have used isometric forces to characterize finger coordination in able-bodied individuals, including enslaving effects (<u>Li et al., 1998</u>) and force-sharing, force-deficit, and enslaving patterns (<u>Zatsiorsky et al., 2000</u>). Similarly, we previously analyzed isometric endpoint forces to characterize impaired finger coordination post-stroke (<u>Xu et al., 2017</u>; <u>2023</u>). However, the spatiotemporal coordination of isometric finger forces post-stroke has not been investigated.

Here, we characterize task-, individual-, and group-level differences in spatiotemporal finger coordination across younger and older able-bodied adults and stroke survivors during isometric force production. We hypothesized that post-stroke neural constraints reduce coordination complexity and expressiveness, resulting in diminished task-, individual-, and group-specificity. We analyzed 3D isometric fingertip forces recorded simultaneously from all five fingers during finger individuation tasks using principal component analysis (PCA) and linear discriminant analysis (LDA). To assess the contributions of salient and subtle features (high- vs. low-variance PCs, respectively), we conducted exploratory analyses of their relative ability in distinguishing tasks, individuals, and groups. We predicted that (1) compared to able-bodied fingers, paretic fingers would require fewer PCs to explain comparable variance in force trajectories, (2) post-stroke coordination would show reduced task- and individual-specificity, (3) paretic fingers would be less group-specific than able-bodied fingers, and (4) high-variance PCs would encode flexor-driven coordination in paretic fingers, while expressiveness would be retained in both high- and low-variance PCs.

## Results

Participants performed finger individuation tasks using the Hand Articulation Neuro-rehabilitation Device (HAND), which stabilizes the hand while independently measuring 3D isometric fingertip forces from each finger (Figure 1A–B). For able-bodied controls and stroke survivors (Supplemental Table 1), fingertip forces were mapped to a virtual coordinate system, allowing participants to control a cursor toward one of six targets corresponding to metacarpophalangeal (MCP) abduction/adduction, proximal interphalangeal (PIP) flexion/extension, and MCP flexion/extension (Figure 1C). Consistent with our prior work characterizing endpoint forces (<u>Xu</u> <u>et al., 2023</u>), paretic fingers generated lower forces with the instructed fingers compared to able-bodied and non-paretic fingers (Figure 1D). Compared to uninstructed able-bodied and non-paretic fingers, uninstructed paretic-finger coactivations were larger (Figure 1E). Phase-aligned isometric force trajectories summed across x-, y-, z- directions qualitatively revealed lower peak force in paretic fingers compared to able-bodied and non-paretic groups (Figure 1F). Angular deviation from the target direction was also greater in paretic fingers (Mean angular deviation = 49.0°) than in younger adult (YA; 41.7°, p = 0.032), older adult (OA; 42.6°, p = 0.007), and non-paretic (NP; 43.2°, p = 0.009) fingers (Figure 1G).

**Figure 1.**
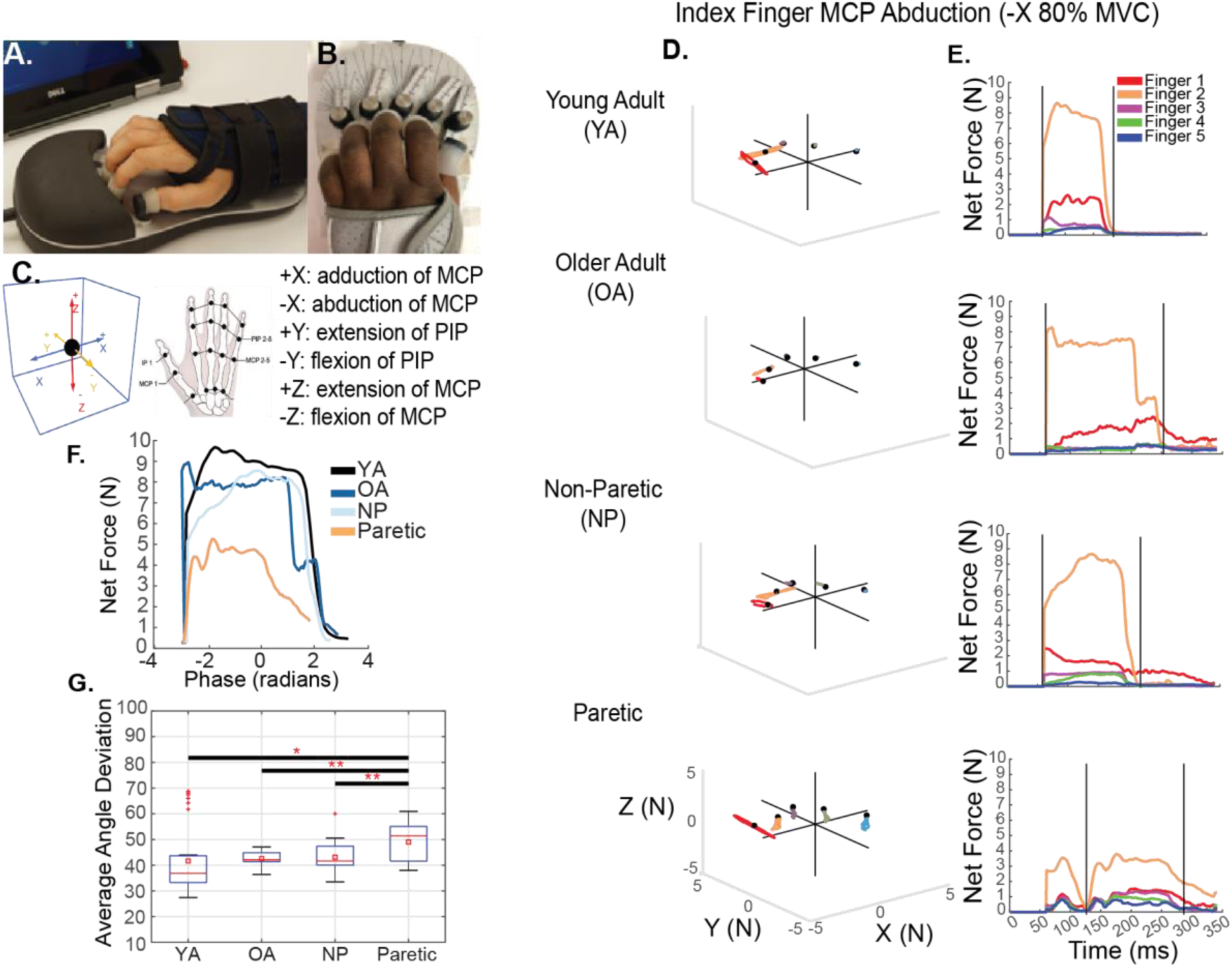
3D isometric force recordings reveal reduced summed force and deviated force vector directions in active fingers post-stroke. **(A-B)** Participants’ hands fitting in the device, hand posture recorded with mounting distance and angle. **(C)** Mapping of isolated force production at different finger joints to Cartesian coordinates in the virtual 3D space. **(D)** Example force trajectories recorded from all fingers of a hand from younger-adult (YA), older-adult (OA), non-paretic (NP), and paretic groups during a finger individuation task (Index Finger Abduction). **(E)** Overall isometric fingertip force trajectories derived from panel D. Vertical black lines illustrate how forces were trimmed in each trial to capture one single force cycle. Greater coactivation of uninstructed fingers and reduced summed force in the instructed finger were observed for post-stroke hands. **(F)** Trimmed data were then phase-averaged to allow comparison of force production from different participants and trials along the same phase range. **(G)** Average angle of deviation between instructed-finger trajectories and the target directions for all groups. Increased deviation angle in post-stroke fingers compared to YA, OA, and NP fingers suggests altered finger coordination in the paretic hand.

### Post-stroke paretic fingers exhibited reduced spatiotemporal coordination complexity compared to non-paretic and able-bodied fingers

Paretic fingers required fewer PCs to explain 95% of the variance accounted for (VAF) in finger forces across tasks or participants, compared to able-bodied and non-paretic fingers (Figure 2A). Paretic fingers required significantly fewer PCs to reach 95% VAF compared with all other groups (Paretic: 8.6 ± 1.6 PCs; NP: 10.5 ± 1.0; OA: 10.7 ± 1.1; YA: 11.1 ± 1.0; all p < 0.010; Figure 2B), whereas no significant differences were observed among the non-paretic fingers and able-bodied groups. PC1 accounted for more of the variance in paretic finger forces (41.0% ± 13.8) than in NP (29.9% ± 7.5), OA (26.3% ± 6.1), and YA (24.8% ± 7.1; all p < 0.001) fingers (Figure 2C). A linear mixed effects (LME) model (fixed effects: Group and PC Number; random effect: Subject) revealed main effects of Group (χ²(3) = 27.6, p < 0.001), PC Number (χ²(14) = 3173.4, p < 0.001), and their interaction (χ²(45) = 638.4, p < 0.001) of the VAF.

**Figure 2.**
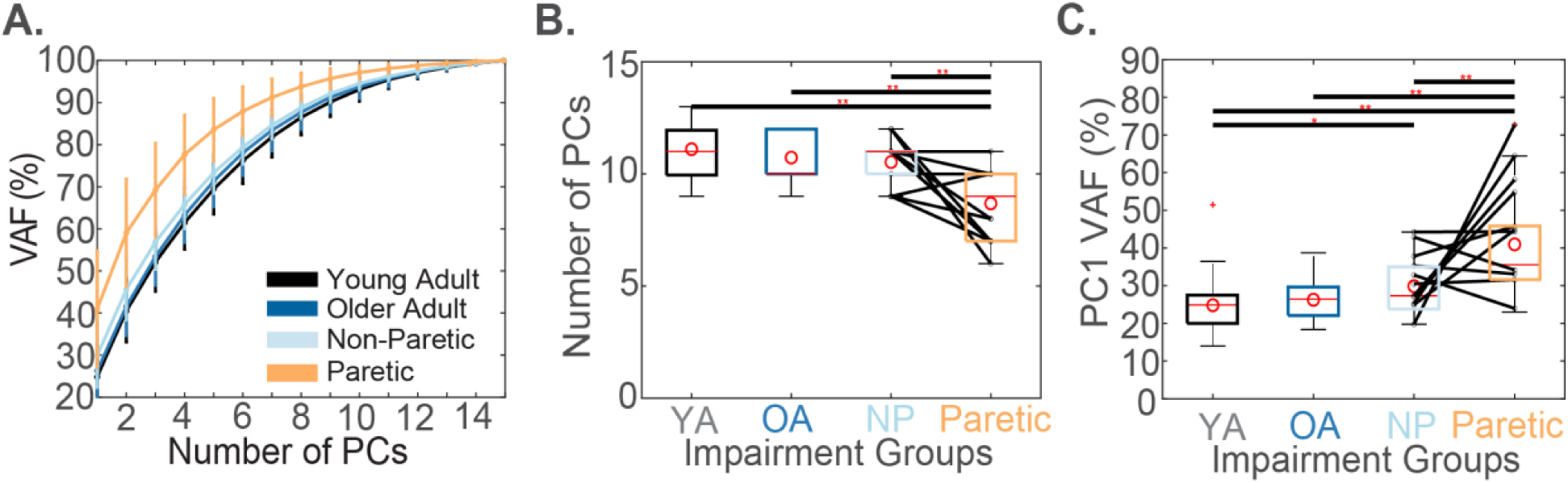
Reduced spatiotemporal finger coordination complexity post-stroke. **(A)** PCA was performed separately for each subject. The cumulative VAF is averaged across the dominant hands of 30 younger-adult (YA), 11 older-adult (OA) able-bodied subjects, and the paretic (Paretic) and non-paretic (NP) hands of 20 stroke survivors. **(B)** Number of PCs needed to account for 95% variance (VAF) and **(C)** PC1 VAF across YA, OA, NP, and Paretic groups. Boxplots show the distribution of PC numbers and PC1 VAF within each group, with red lines and circles indicating the median and mean values, respectively. Paretic hands require significantly fewer PCs and paretic PC1 accounts for significantly more variance compared to YA, OA, and NP hands, suggesting reduced complexity in the paretic hand post-stroke.

### High- and low-variance PC trajectories differed qualitatively between target fingers and tasks

Qualitative inspections of high- (PCs 1-3) and low-variance (PCs 13-15) PC activations revealed distinct coordination patterns across groups and finger individuation tasks (instructed fingers and target directions). High-variance PC activations appeared to differ across instructed fingers (left plots in Figure 3A–D; the same task is shown for two instructed fingers) and target directions (left plots in Figure 3E–H; the same finger is shown for two tasks). This task-specific activation separation persisted even in low-variance PCs (right plots in Figure 3E-H). PC activations varied across fingers and tasks in all groups (example participants from YA, OA, NP, and Paretic shown in Figure 3).

**Figure 3.**
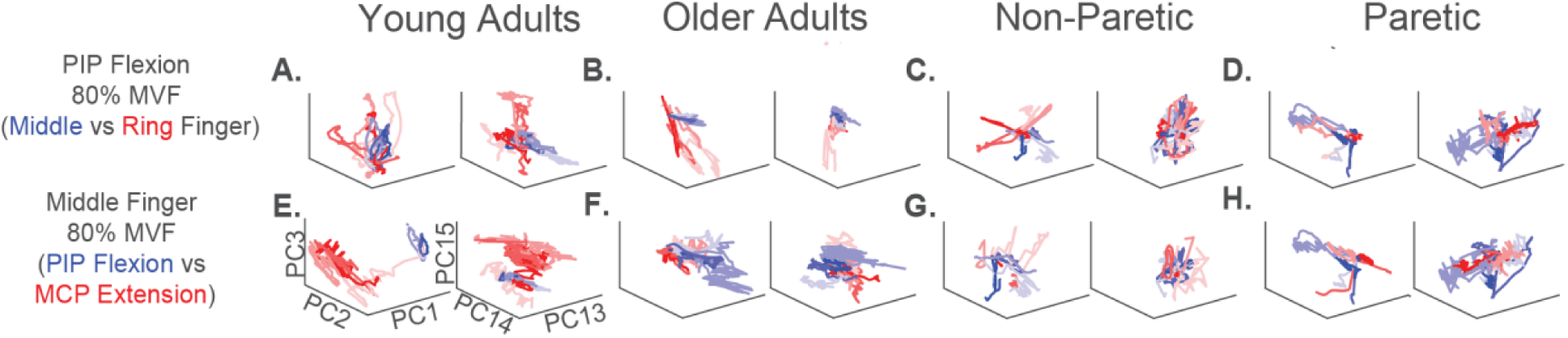
High- and low- variance PC projections of able-bodied and post-stroke force trajectories across different task. Time-varying PC coactivations are shown for a younger-adult and older-adult able-bodied individual (dominant hand) and a stroke survivor (non-paretic and paretic hands) performing different finger force production tasks. Each row highlights differences in PC coactivations between two task conditions—plotted in red and blue—for able-bodied, non-paretic, and paretic hands. Comparisons include: **(A–D)** finger comparison: middle vs. ring finger PIP flexion at 80% MVF, and **(E–H)** joint comparison: middle finger PIP flexion vs. MCP extension at 80% MVF. Four repeated trials per condition are shown in varying shades. PC coactivations for each task condition are plotted in red and blue, with four repeated trials in varying shades. The left panels in each column show projections onto high-variance PCs (PCs 1–3), while the right panels show projections onto low-variance PCs (PCs 13–15). In both cases, PC coactivations for different tasks occupy distinct regions, indicating that low-variance PCs do not merely capture noise but also encode structured task-specific information, such as temporal and spatial coordination patterns.

### Paretic fingers presented reduced task- and group-specificity but comparable individual-specificity to non-paretic and able-bodied older adult fingers

When classifying tasks using linear discriminant analysis (LDA) and a component space consisting of all 15 trial-averaged PCs, the ability to classify tasks was lower in paretic fingers (mean accuracy = 66.4% ± 28.6) compared to NP (accuracy = 90.4% ± 6.0), OA (accuracy = 92.4% ± 6.1) and YA (accuracy = 97.1% ± 3.6; all p < 0.001; Figure 4A,G) fingers. The variance in classification accuracy for paretic fingers was larger compared to other groups (e.g., ± 28.6 vs 6.0% in Paretic vs NP fingers). The ability to classify individuals using all PCs was also the highest in YA (accuracy = 86.6% ± 7.7) compared to OA (accuracy = 79.3% ± 8.3; p < 0.001), NP (accuracy = 77.3% ± 8.3; p < 0.001), and Paretic (accuracy = 78.2% ± 8.3; p < 0.001) fingers. In paretic fingers, greater intrusion of a flexor bias classification was associated with classification accuracy for tasks (r = −0.83, p < 0.001; Figure 4D), individuals (r = −0.48, p = 0.036; Figure 4E), and groups (r = -0.72, p < 0.001; Figure 4F). The ability to classify groups using all PCs was lower for paretic fingers (mean accuracy = 53.8% ± 10.0) compared to NP (accuracy = 54.7% ± 10.0; p = 0.046), OA (accuracy = 61.2% ± 10.4; p < 0.001), and YA (accuracy = 70.0% ± 12.3; p < 0.001) fingers (Figure 4C,I). LME models (fixed effects: Group and PC Removal Number; random effect: Subject) identified main effects of Group (χ² > 43.1, p < 0.001), PC Removal Number (χ² > 1364.2, p < 0.001), and their interaction (χ² > 163.8, p < 0.001) on task, individual, and group classification accuracy.

**Figure 4:**
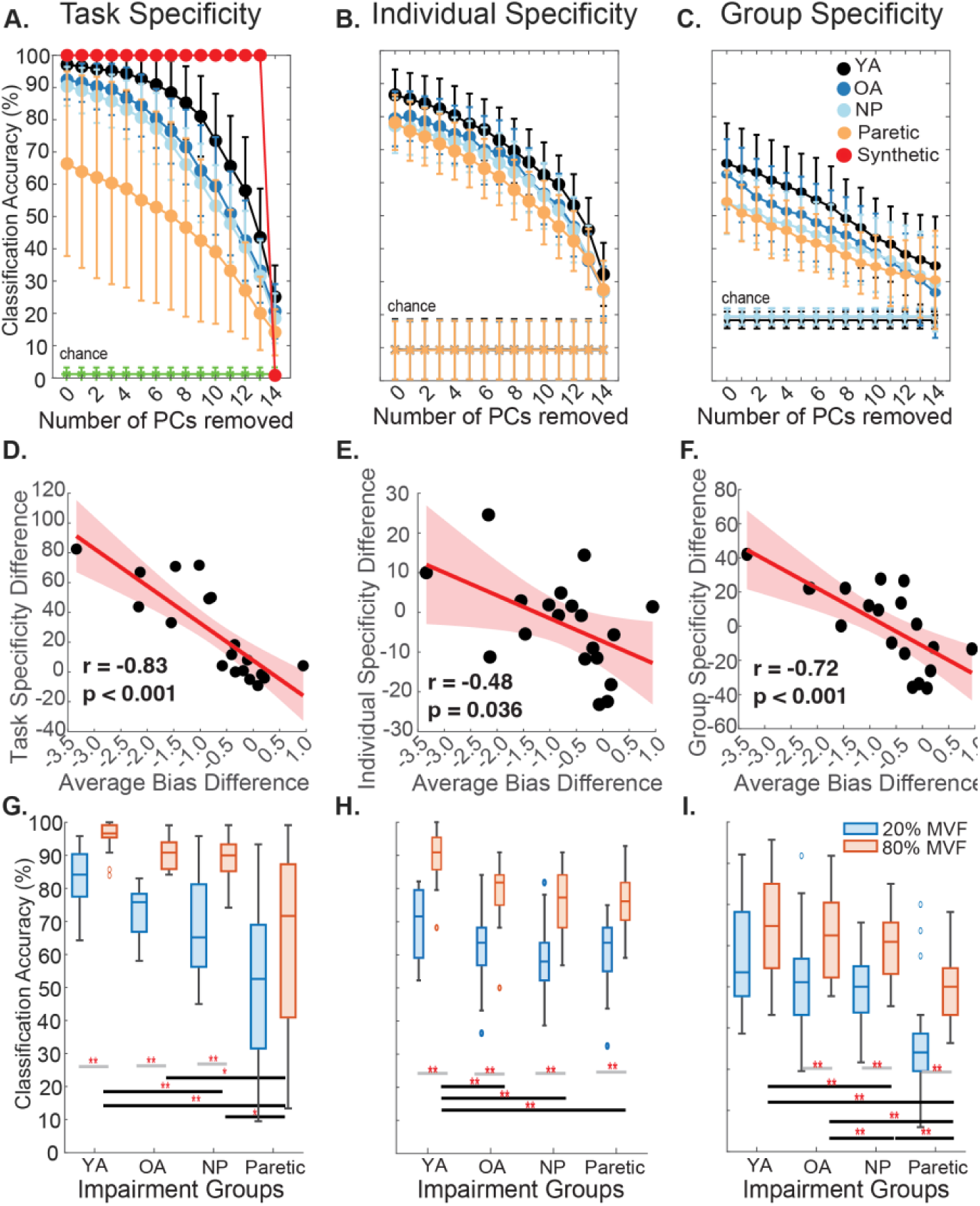
Reduced task-, individual- and group-specificity in post-stroke finger coordination. PCA was applied to 3D isometric fingertip force data collected during finger individuation tasks. Time-varying principal component (PC) coactivations, concatenated for each participant performing across all tasks, were used as input for an LDA classifier to distinguish between different tasks **(A,D,G)**; those concatenated across all participants performing the same task were used for classifying individuals **(B,E,H)** and groups **(C,F,I)**. Classification accuracy for **(A)** tasks, **(B)** individuals, and **(C)** groups was further evaluated at 80% MVF after progressively removing high-variance PCs. **(A)** The red curve shows task classification accuracy obtained from a simulated dataset containing randomly generated signals with additive Gaussian noise, processed with the same procedure as the experimental data (temporal PC-removal and LDA) to evaluate whether sensor noise alone could produce above-chance classification from lower-variance components (Supplemental Figure 1). **(C, F)** Scatter plots show that reduced task- (r = –0.83, p < 0.001), individual- (r = –0.48, p = 0.036), and group-specificity (r = –0.72, p = 0.001) in paretic hands are correlated with post-stroke intrusion of flexor bias (Xu et al., 2023). **(G, H, I)** Task-specific **(G)**, individual-specific **(H)**, and group-specific **(I)** classification accuracies across tasks were compared at 20% (orange) and 80% (blue) of Maximum Voluntary Force (MVF).

As expected, iteratively removing high-variance PCs reduced the ability to classify tasks, individuals, and groups using finger force coordination (Figure 4A-C, respectively). Even when classifying using only the lowest-variance PC as features, LDA classification accuracy for tasks was greater than chance for all groups (mean PC15 classification accuracy for tasks; YA = 25.2% ± 9.7%; OA = 20.7% ± 8.4%; NP = 18.5% ± 7.7%; Paretic = 14.2% ± 7.2%; all p < 0.001; Figure 4A). Classification accuracy for individuals (mean PC15 classification accuracy for individuals; YA = 31.0% ± 8.9%; OA = 27.1% ± 7.5%; NP = 26.9% ± 8.5%; Paretic = 27.0% ± 9.2%; all p < 0.001; Figure 4B) and groups (mean PC15 classification accuracy for groups; YA = 25.2% ± 9.7%; OA = 20.7% ± 8.4%; NP = 18.5% ± 7.7%; Paretic = 14.2% ± 7.2%; all p < 0.001; Figure 4C) were greater than chance for all groups. Analyses of synthetic data confirmed that classifications using PC15 did not reflect mathematical artifacts of PCA decomposition, as the lowest-variance components derived from simulated data did not retain above-chance classification accuracy (Figure 4A; Supplemental Figure 1).

All classifications using reduced PC models (Figure 4A-C) used data from trials where participants reached 80% of their maximum voluntary force (MVF) levels—the highest force level tested—during each task. We focused our analyses on the 80% MVF trials because this higher force level resulted in greater task, individual, and group classification accuracy than lower MVF targets. For example, the 80% MVF trials exhibited stronger task (p < 0.001), individual (p < 0.001), and group (p < 0.001) classification accuracy than the same tasks performed at 20% MVF (Figure 4G-I).

### Higher-variance PCs tended to contribute more to task and group-specificity, while both high- and low-variance PCs contributed to individual-specificity equally

Across tasks, LDA coefficients for different PCs were consistently non-zero across group pairs (Figure 5A; example shown for little finger MCP adduction; rows denote group pairs), with asterisks denoting components that differed significantly from zero across participants, indicating reliable contributions to group separation. For example, PC1 contributed significantly when classifying paretic and older adult fingers versus younger adult fingers, , but not in other comparisons. To interpret the coordination patterns captured by these components, we reconstructed the significantly contributing PCs in 3D force space (Figure 5B, Figure 6; see a complete catalog in Supplemental Figure 2).

**Figure 5:**
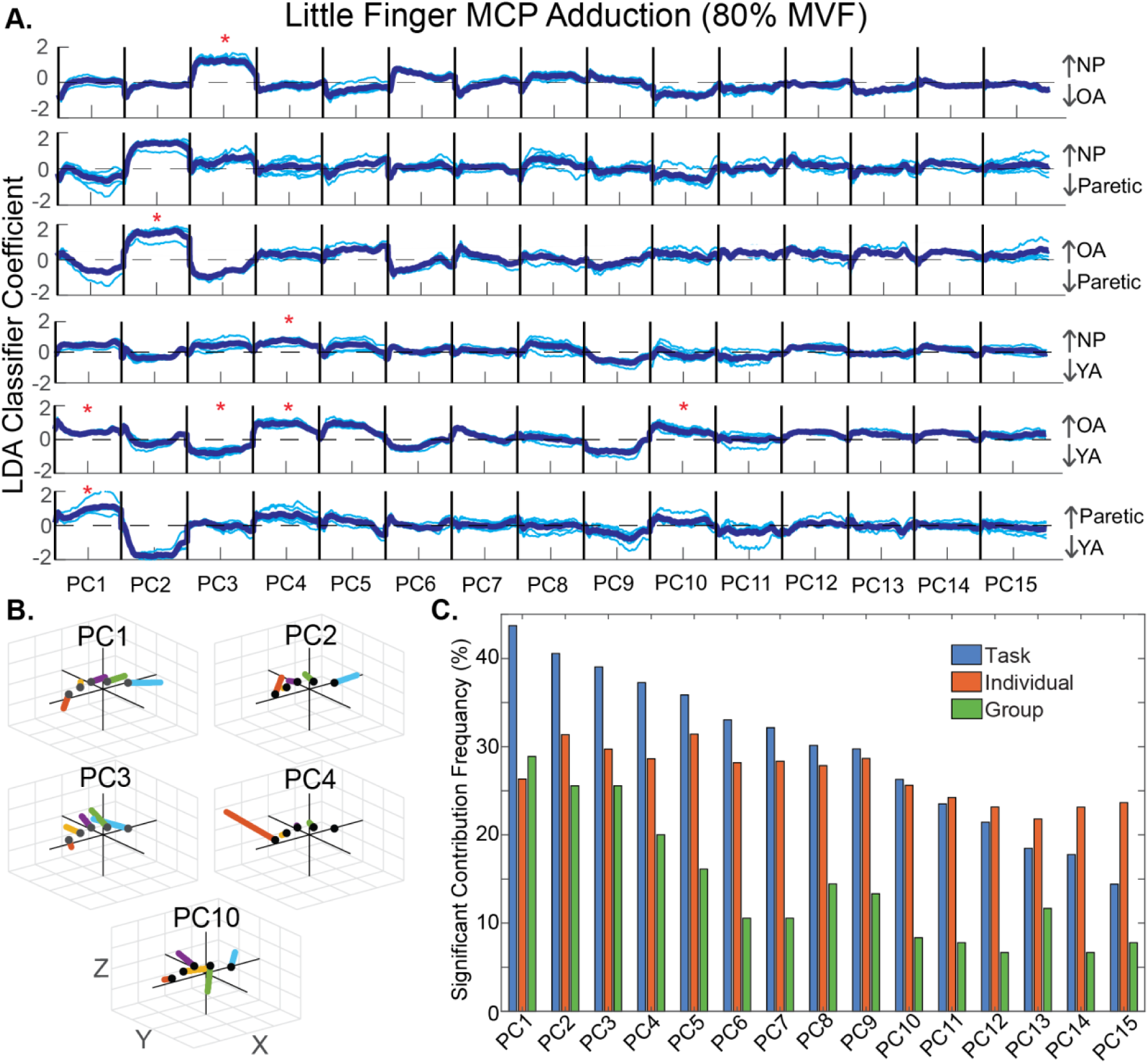
LDA coefficients reveal the most diagnostic PCs for identifying differences in coordination across tasks, individuals, and groups. **(A)** Representative example of Z-score normalized, time-varying PC coactivations from finger force tasks were used in pairwise group LDA classifications. Each PC spans 0–100% of the normalized force trajectory. LDA coefficients indicate each PC segment’s contribution to group discrimination, with positive values favoring Class 1 and negative values favoring Class 2. Statistical PC contribution significance was determined by bootstrapping 99% confidence intervals (CIs); asterisks mark the segments whose coefficients significantly contributed to pairwise classification. Light blue lines show coefficients from all bootstrap iterations; thicker dark blue lines show the average across iterations. **(B)** Representative examples of the 3D vector representation of PC coefficients3D vector representations of PC coefficients identified as significantly contributing to LDA classification are presented in 3D. **(C)** Bar graph showing the frequency with which individual PCs significantly contributed to distinguishing between different tasks (blue), individuals (orange), and groups (green).

**Figure 6:**
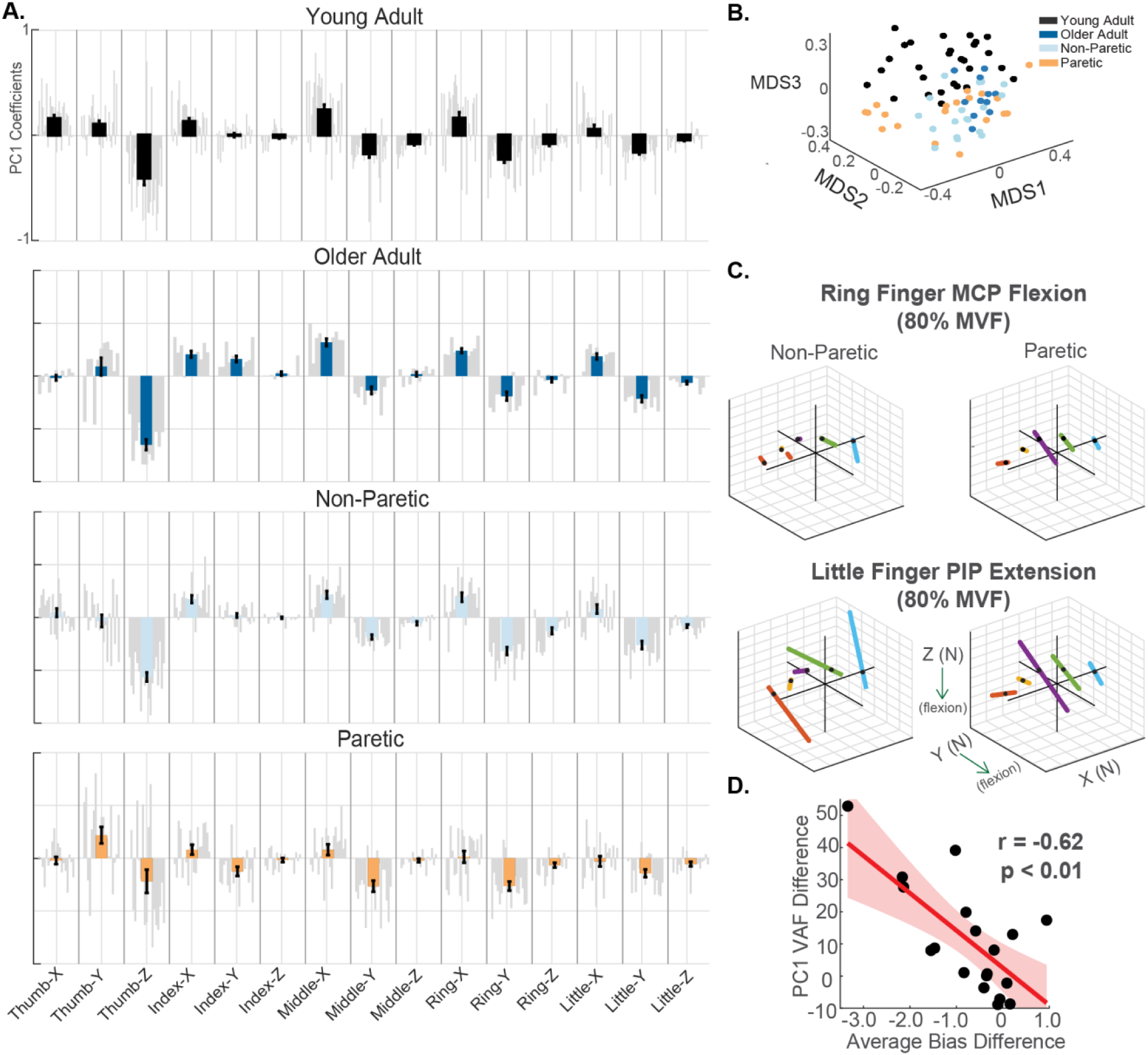
Altered PC1 components correlate with severity in flexor bias intrusion in paretic hands. **(A)** PC1 coefficients across 15 finger-axis force channels for each group: younger adults, older adults, non-paretic, paretic hands. Gray bar represents an individual’s PC1 loading per channel; colored overlays denote group means. Paretic hands show a bias toward flexion-direction channels. **(B)** 3D multidimensional scaling (MDS) embeddings of pairwise PC1 variance accounted for (VAF) reveal similarities in PC1 components across individuals with similar impairment conditions. **(C)** PC1 was used to reconstruct isometric fingertip force trajectories for non-paretic and paretic hands of an exemplar stroke survivor during Ring-finger Flexion at the MCP joint and Little-finger Extension at the PIP joint at 80% MVF. **(D)** The paretic vs. non-paretic difference in PC1 variance was negatively correlated (r = −0.62, p < 0.01) with the intrusion of flexor bias.

Higher-variance PCs (PCs 1–7) contributed more often than lower-variance PCs (PCs 8–15) to task- (χ²(1) = 12532.3, p < 0.001, Cramer’s V = 0.16) and group-specific (χ²(1) = 33.5, p < 0.041, Cramer’s V=0.11) classifications, indicating small but consistent effects. For individual-specificity, high- and low-variance PCs contributed to classification accuracy statistically differently (χ²(1) = 2017.6, p < 0.001) though the magnitude of the difference was negligible (Cramer’s V = 0.05). The contributions were more evenly distributed across PCs (27.9% in higher-variance PCs vs. 23.9% in lower-variance PCs; Figure 5C).

### High- and low-variance PCs captured different aspects of spatiotemporal finger coordination

PC1 coefficient magnitudes, where positive and negative loadings reflect flexor and extensor contributions, respectively, revealed stronger flexor-dominated components across paretic fingers compared to able-bodied and non-paretic fingers (Figure 6A). PC1 coefficients were highly consistent across subjects within each group, in contrast to the greater inter-individual variability observed in lower-variance components (PC15; Figure 7A). PC1 coefficients differed between paretic and both non-paretic and able-bodied fingers, reflecting altered coordination patterns post-stroke. To visualize these between-group differences, we applied multidimensional scaling (MDS) to individual PC1 coefficient vectors. In this embedding, each point represents one participant, and the distance between points reflects the similarity of their PC1 coefficient patterns, such that participants with more similar patterns cluster closer together in the reduced-dimensional space (Figure 6B). Reconstructions of 3D Isometric fingertip forces using only PC1 demonstrated that in paretic fingers, both flexion and non-flexion tasks collapsed into trajectories dominated by flexion patterns in the non-instructed fingers, as illustrated by an 80% MVF (Figure 6C; ring-finger MCP flexion task and little-finger PIP extension task shown as two examples). This observation is supported by a negative correlation between the variance explained by PC1 and the magnitude of flexor bias intrusion across participants (r = −0.62, p < 0.001; Figure 6D).

**Figure 7:**
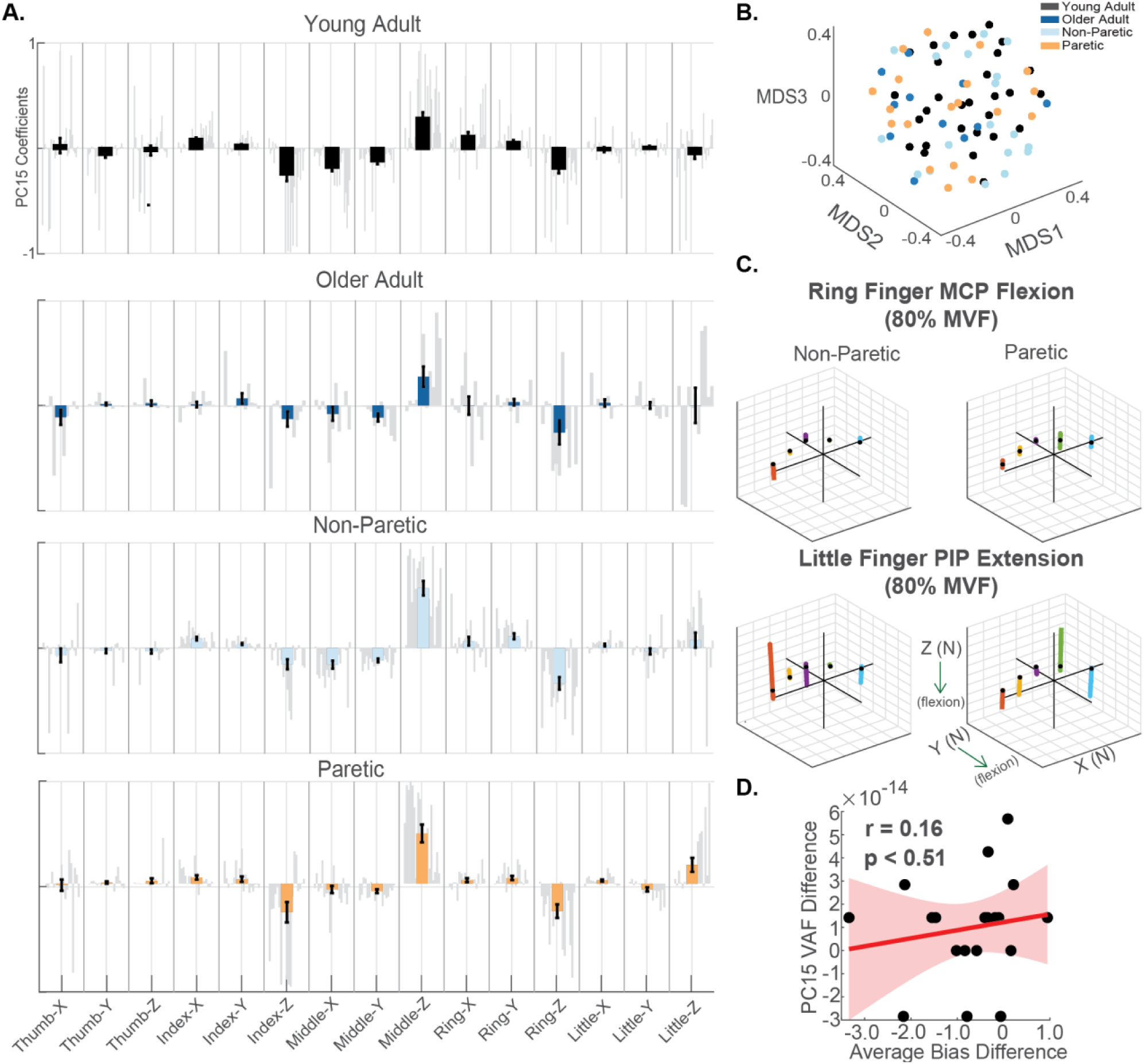
Less stereotyped coordination patterns characterize lower-variance PCs. **(A)** PC15 coefficients across 15 finger-axis force channels for each group: younger adults, older adults, non-paretic, paretic hands. Gray bar represents an individual’s PC15 loading per channel; colored overlays denote group means. **(B)** 3D multidimensional scaling (MDS) embeddings of pairwise PC15 variance accounted for (VAF) illustrate high individual variability across all groups, suggesting less stereotyped coordination patterns. **(C)** PC15 was used to reconstruct isometric fingertip force trajectories for non-paretic and paretic hands of an exemplar stroke survivor during Ring-finger Flexion at the MCP joint and Little-finger Extension at the PIP joint at 80% MVF. **(D)** The paretic vs. non-paretic difference in PC15 variance was not correlated (r = 0.16, p = 0.61) with the difference in force bias averaged across the Y and Z directions.

Unlike PC1, whose coefficients were similar within groups and systematically shifted in paretic fingers, PC15 coefficients appeared heterogeneous both within and across groups. Consistent with this observed variability, an MDS projection of PC15 coefficients did not reveal clear clustering by group (Figure 7B). Reconstructions using only PC15 did not produce flexor-dominated patterns (Figure 7C), and the variance explained by PC15 was not correlated with the magnitude of flexor bias intrusion (r = 0.16, p = 0.513; Figure 7D), suggesting that PC15 does not capture the post-stroke intrusion of flexor bias observed in paretic fingers.

To test whether the above-chance classification based on low-variance PCs was due to mathematical artifacts, we generated synthetic 14-dimensional signals for 120 task conditions across noise levels (approximately 0%, 1.0%, 2.0%, and 5.0% of the maximum signal amplitude; Supplemental Figure 1) and projected them onto 15 dimensions, such that PC15 contained only noise. When PCA was applied to synthetic force trajectories, the lowest-variance components did not exhibit consistent above-chance classification performance.

### Compared to endpoints, spatiotemporal finger coordination patterns showed greater classification accuracy for tasks, individuals, and groups

We conducted the same PCA and LDA analyses over endpoint data computed as all fingertip forces when the instructed finger reaches its peak force. LME models were then used to compare LDA classification accuracies across Measurement Types (endpoint vs. spatiotemporal). For task specificity, the LME model revealed a significant main effect of PC Removal Number (χ²(14) = 777.2, p < 0.001), but no effect of Measure Type (χ²(1) = 1.7, p = 0.190) or interaction (χ²(15) = 22.3, p = 0.074). Task classification accuracy declined as high-variance PCs were removed, but did not differ between spatiotemporal and endpoint measures. When using all PCs, classification accuracy was comparable between measures. Descriptively, when classification relied only on the lowest-variance PC (PC15), spatiotemporal data showed slightly higher accuracy than endpoints across groups (mean difference between spatiotemporal and endpoint measures: YA: 9.6%; OA: 7.0%; NP: 7.0%; Paretic: 4.6%; Figure 8A–D), with 95% confidence intervals overlapping between measure types, consistent with the absence of a significant main effect of Measure Type. For individual specificity, the LME model revealed significant main effects of Measure Type (χ²(1) = 4.9, p = 0.023) and PC Removal Number (χ²(14) = 2402.0, p < 0.001), and their interaction (χ²(15) = 772.0, p < 0.001). Spatiotemporal coordination patterns classified individuals more accurately than endpoint patterns, and this advantage depended on the number of PCs retained. Across PC sets, spatiotemporal data consistently outperformed endpoint data in YA, OA, NP, and paretic fingers (Figure 8E–H). For group specificity, a significant Measure Type × PC Removal Number interaction was observed (χ²(15) = 34.3, p < 0.005; Figure 8). Spatiotemporal patterns showed higher classification accuracy than endpoint patterns when all PCs were retained (mean difference between the two measures: YA: 6.7%; OA: 3.5%; NP: 2.7%; Paretic: 4.7%), but not when classification relied only on low-variance PCs (Figure 8I–L).

**Figure 8:**
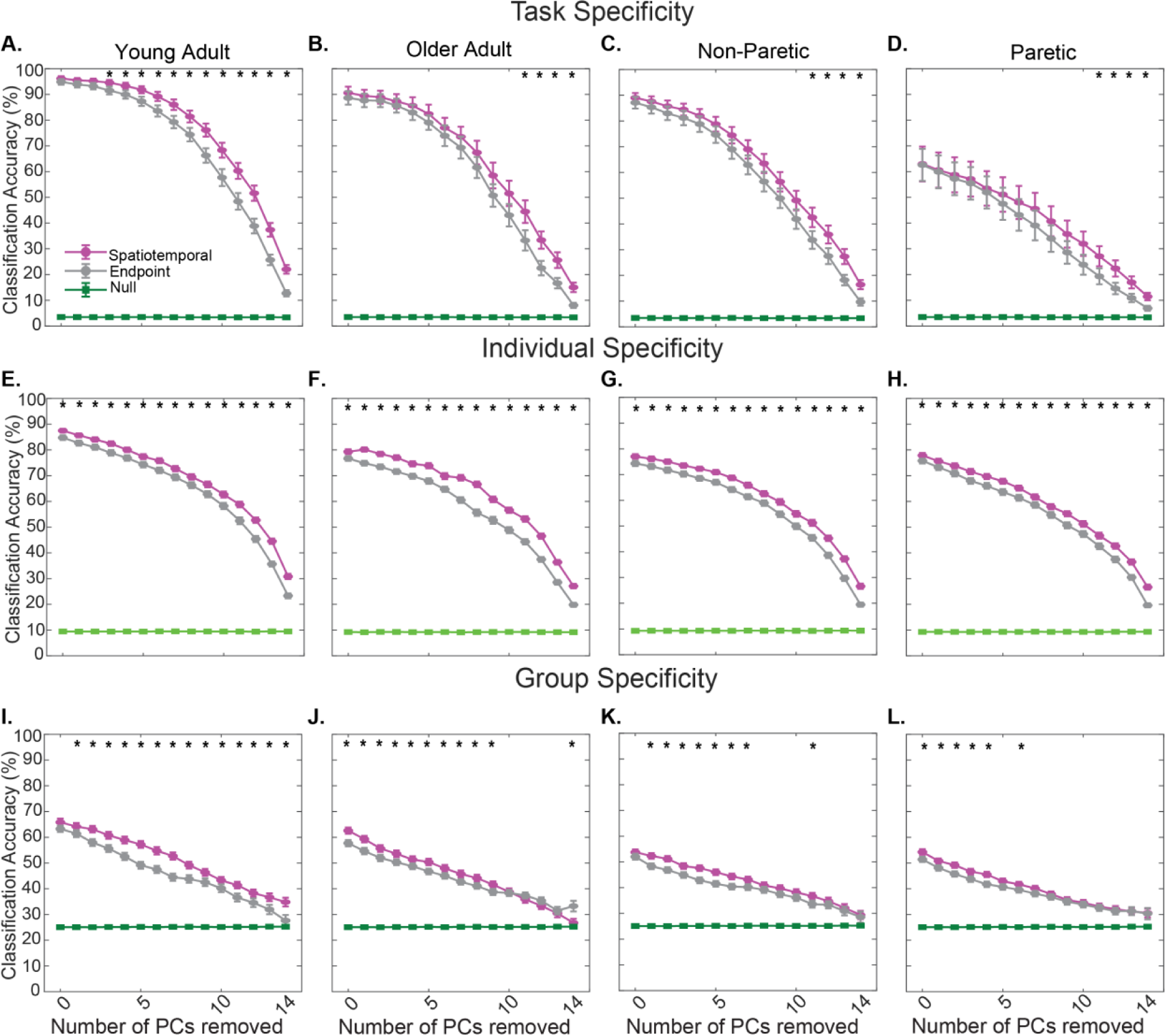
Task-, individual-, and group-specific classification accuracies for endpoint (gray) and spatiotemporal (purple) forces across impairment groups. Statistical significance was assessed by comparing 95% confidence intervals between endpoint and spatiotemporal accuracies for each principal component (PC). Black asterisks mark PCs with non-overlapping 95% confidence intervals (approximate p < 0.050). Across all impairment conditions, spatiotemporal forces consistently yielded higher task- and individual-specific classification accuracies than endpoint forces, and high-variance spatiotemporal PCs captured greater group-level variability.

## Discussion

This study characterized 3D spatiotemporal force coordination patterns across all five fingers in able-bodied participants and chronic stroke survivors. Stroke survivors exhibited reduced coordination complexity (fewer PCs), and reduced expressiveness (task- and individual-specificity) compared to able-bodied adults. Reductions in expressiveness were associated with more severe intrusion of flexor bias in paretic fingers. Compared to endpoint forces, spatiotemporal coordination showed slightly greater task and individual expressiveness, and group specificity, indicating that temporal aspects of finger coordination contain meaningful information about tasks, individuals, and groups. Exploratory analyses revealed that both salient and subtle aspects of coordination (i.e., high- and low-variance PCs, respectively) contributed to individual expressiveness, whereas more salient components captured task- and group-differences. Collectively, these findings suggest that post-stroke neural constraints reduce coordination complexity and expressiveness, which may reduce stroke survivors’ abilities to coordinate finger forces during diverse activities of daily living and may represent useful targets for post-stroke hand rehabilitation.

### Stroke survivors exhibit reduced spatiotemporal finger coordination complexity

Specifically, paretic finger coordination is less complex than that in both able-bodied and non-paretic fingers. This finding is consistent with prior studies of endpoint finger coordination, which demonstrated increased inter-finger coupling during individuated finger force production tasks (<u>Lang & Schieber, 2003</u>, <u>2004</u>; <u>Li et al., 2003</u>; <u>Xu et al., 2017</u>). Reduced coordination complexity in paretic fingers may emerge from stroke-related neural and biomechanical constraints. This interpretation is consistent with findings of reduced complexity in post-stroke muscle coordination during walking (<u>Cheung et al., 2009; Clark et al., 2010</u>), upper limb movements (<u>Roh et al., 2013</u>, <u>2015</u>), and isometric fingertip force production (<u>Xu et al., 2023</u>). During isometric tasks, altered neural constraints on muscle coordination may limit the feasible set of coordination patterns available to produce the target forces (<u>Valero-Cuevas et al., 1998</u>). Muscle activity was not recorded in the present study, such that future studies should investigate how this reduced coordination complexity reflects altered neural constraints on post-stroke muscle coordination.

### Task- and individual-expressiveness and group-specificity are reduced in post-stroke paretic fingers

High classification accuracy for tasks, individuals, and groups from coordination patterns suggests that the spatiotemporal characteristics of finger forces encode information about the task being performed, who is performing it, and whether the fingers are able-bodied (YA or OA), non-paretic, or paretic. This observation aligns with prior work showing that task information is encoded in coordination components, including in low-variance PCs (<u>Yan et al., 2020</u>). However, task discrimination was more variable in paretic fingers than able-bodied or non-paretic fingers. One possible explanation for this variability is the intrusion of flexor bias, which constrains the coordination patterns available to paretic fingers. Consistent with this hypothesis, task-specificity decreased as the intrusion of flexor bias increased. These findings indicate that stroke-related flexor coupling may reduce the separability of task-specific coordination. These findings extend prior findings linking flexor bias intrusion with increased finger coactivation magnitude across tasks (<u>Xu et al., 2023</u>).

Higher force levels enhanced coordination expressiveness, producing more distinct spatiotemporal patterns across tasks, individuals, and groups. This finding likely reflects increased motor unit recruitment (<u>Fuglevand et al., 1993</u>) and stronger finger coupling at higher forces (<u>Valero-Cuevas, 2000</u>), consistent with prior findings that post-stroke impairments in individuation and inter-finger coupling become more pronounced at higher force levels (<u>Li et al.,</u> <u>2003</u>; <u>Xu et al., 2017</u>). Thus, clinically, higher-force tasks may improve sensitivity to coordination deficits, though this effect should be interpreted cautiously, as increased force may also increase variability and exaggerate observable differences in coordination (e.g., <u>Fuglevand</u> <u>et al., 1993</u>; <u>Slifkin & Newell, 1999</u>). Notably, the highest force tested (ten Newtons), may still be below the maximum motor output for the able-bodied and mildly impaired participants.

Consistent with prior findings using postural endpoint forces in able-bodied adults (<u>Yan</u> <u>et al., 2020</u>), even the most subtle components of post-stroke finger force coordination (i.e., the lowest-variance PCs) in the current study encoded information about tasks, individuals, and groups. One possible explanation is that low-variance PCs capture stylistic or idiosyncratic coordination that varies across individuals and are only weakly constrained by task demands, as shown by the wide spread in the MDS plot (Fig. 8B). Although above-chance classification accuracy of the lowest-variance PCs may be an artifact of PCA applied to noisy measurements or experimental biases introduced during setup or sensor zeroing. However, synthetic data simulations showed that applying PCA to noisy signals alone did not produce above-chance classification accuracy, suggesting that noise is unlikely to account for this effect. We also did not identify a source of bias that would explain the above-chance classification accuracy for tasks and individuals. Together, these findings suggest that low-variance PCs capture aspects of coordination rather than measurement noise or bias, potentially capturing subtle, task- and individual-specific aspects of coordination that contribute to the expressive repertoire in both able-bodied and post-stroke finger forces. We explored how subtle components of coordination contribute to task-, individual-, and group-specificity, and how these components differ from more-salient components in their relationship to task constraints and impairment.

### Task-Specificity

For able-bodied and post-stroke fingers, task-specific information is primarily encoded in salient components, though subtle components also retain task-relevant information. This observation suggests that task identity is distributed across both high and low-variance components, a trend also preserved in paretic fingers despite reduced differentiation of coordination patterns.

Our findings also extend prior evidence that subtle components capture meaningful aspects of coordination in able-bodied fingers. Whereas previous work showed that subtle coordination components can distinguish grasp types and sign language gestures in able-bodied fingers (<u>Yan et al., 2020</u>) and task parameters in musculoskeletal hand models (<u>Chiappa</u> <u>et al., 2024</u>), the task-specificity of subtle aspects of coordination generalize to spatiotemporal finger force coordination in the current study, even in post-stroke paretic fingers. Unlike more salient components, subtle components of paretic finger coordination are not related to the intrusion of flexor bias, consistent with our previous finding that reduced geometric complexity in endpoint finger coordination — defined as reduced task-dependent variance in the geometric shape of coordination patterns— was independent of flexor-bias intrusion (<u>Xu et al., 2023</u>). Salient and subtle components of coordination may also reflect biomechanical differences that influence finger coordination, such as strength differences across fingers, joint coupling, and tendon biases that would be differentially challenged across tasks (<u>Jarrassé et al., 2014</u>). However, because we used an isometric force production task that mitigates the effects of time-varying musculotendon states, salient and subtle components of coordination observed in our data likely mostly capture differences in neural control of finger coordination.

### Individual-Specificity

Our findings suggest that both salient and subtle components of coordination carry distinct information about each person’s unique neural control strategy, even after a stroke. This interpretation is consistent with prior studies showing that individual expressiveness in motor coordination is evident across a range of behaviors, including spatiotemporal coordination of walking (<u>Winner et al., 2023</u>) and culturally shaped manual skills such as pottery (<u>Gandon et al., 2013</u>). Further, our interpretation aligns with the broader view that human motor coordination reflects personalized control patterns sculpted by lifelong learning and experience (<u>Ting et al., 2024</u>).

### Group-Specificity

One possible interpretation of reduced group-specificity in paretic fingers compared to able-bodied fingers is that stroke does not produce a single, consistent pattern of multi-finger coordination within the paretic group. While general neural and biomechanical characteristics are observed post-stroke, such as intrusion of flexor bias (<u>Xu et</u> <u>al., 2023</u>), differences in lesion locations and compensatory strategies may lead to heterogeneous finger coordination across stroke survivors, weakening consistent group-level characteristics. Consequently, a single, well-defined pattern of stroke-impaired finger coordination may not exist. This interpretation aligns with prior work demonstrating heterogeneous muscle coordination during walking (<u>Cheung et al., 2012</u>) and in upper-limb force control after stroke (<u>Lodha et al., 2010</u>, <u>2013</u>), which may be associated with differences in lesion characteristics or compensatory strategies and movement patterns.

### While salient components primarily drive task- and group-specific classification, both salient and subtle components retain individual expressiveness

Our exploratory analysis of LDA coefficient analysis suggests that salient components of coordination (i.e., higher-variance PCs) are more characteristic of global changes in coordination from task demands and impairment constraints. These findings suggest salient features may reflect force coordination patterns shaped by task constraints and coarse impairment-related characteristics (e.g., intrusion of flexor bias). This interpretation is consistent with prior work showing that finger coordination is strongly shaped by task goals and impairment-related biomechanical limitations (<u>Santello et al., 1998</u>; <u>Ingram et al., 2008</u>; <u>Lang &</u> <u>Schieber, 2003</u>).

In contrast, both salient and subtle coordination components appear characteristic of individual expressiveness. Qualitative comparisons of salient coordination components revealed more differentiated coordination across fingers and target directions in able-bodied and non-paretic fingers, whereas paretic fingers exhibited flexion-biased patterns, consistent with well-documented flexor-dominant coupling post-stroke (<u>Dewald et al., 1995</u>; <u>Lodha et al., 2010</u>). Subtle components, in contrast, revealed heterogeneous patterns across fingers and target directions compared with salient components. Reconstruction of 3D fingertip force trajectories using PC15 structures further illustrated these patterns. Consistent with this observation, salient components of paretic finger coordination were more strongly associated with the intrusion of flexor bias compared to subtle components of paretic finger coordination. Multidimensional Scaling of salient components showed partial separation between groups, with paretic fingers exhibiting greater dispersion than able-bodied and non-paretic fingers, whereas that for subtle components revealed greater within-group variation for all groups compared to salient components. Together, these findings indicate that salient features reflect coordination patterns shaped by task constraints and coarse impairment-related characteristics, while finer variations in individual expressiveness are distributed across salient and subtle components.

### Compared to endpoints, spatiotemporal finger forces suggest similar levels of coordination complexity, but more characteristics of tasks, individuals, and groups

Prior work has predominantly characterized finger coordination complexity using endpoint postures or static forces (<u>Santello et al., 1998</u>; <u>Santello & Soechting, 2000</u>; <u>Jarque-Bou et al.,</u> <u>2019</u>). Here, we demonstrated that differences in coordination complexity between able-bodied adults and stroke survivors are similar when computed using endpoint or spatiotemporal forces. However, spatiotemporal patterns encoded slightly more information about tasks, individuals, and groups than endpoint forces, reflected in greater classification accuracy (1.7-9.7% greater) when using spatiotemporal forces. The slightly greater expressiveness of spatiotemporal forces is likely because they capture individual differences in *how* force trajectories *evolve* over time, even within tasks where similar endpoint force magnitudes are produced. For example, similar endpoint measures in speed, accuracy, or smoothness during pointing can be achieved using distinct coordination strategies (<u>Levin et al., 1996</u>; <u>Levin et al., 2002</u>; <u>van Kordelaar et al., 2013</u>). Although it remains to be investigated whether this additional expressiveness is clinically meaningful, we speculate that spatiotemporal information may reveal aspects of impaired coordination not apparent from endpoints alone. Future work should examine whether distinct spatiotemporal coordination phenotypes relate to functional outcomes post-stroke.

### Limitations

Several factors may limit the generalizability of our findings. First, tasks were restricted to isometric forces. While mitigating biomechanical confounders due to motion, this paradigm does not capture the richness of naturalistic finger use that involves non-isometric tasks and more complex variations in kinematics and musculotendon states (<u>Ingram et al., 2008</u>; <u>Valero-Cuevas</u> <u>et al., 2009</u>). Future work considering control of non-isometric finger motion may be useful in characterizing spatiotemporal coordination patterns in more ecologically valid settings. Second, PCA is limited to identifying linear structures; consequently, aspects of coordination that emerge through nonlinear interactions may not be fully captured. Future work using nonlinear dimensionality-reduction techniques, such as autoencoders and recurrent neural networks, could help determine whether coordination is organized in nonlinear manifolds (<u>Nathella et al.,</u> <u>2025</u>) and whether such coordination reflects similar or further reduced complexity. Finally, although we observed effects of stroke in primary between-group comparisons, it remains unclear whether distinct clinical subtypes exhibit systematically different spatiotemporal coordination patterns.

## Conclusion

Here, we demonstrated that stroke survivors exhibit reduced complexity and expressiveness of multi-finger isometric force coordination, suggesting that they have a limited repertoire of coordination patterns available for finger force production. Characteristics of post-stroke impairments and task constraints appear to be reflected in the more salient components of finger coordination. However, stroke-impaired fingers retain task-, individual, and group-specific information in both salient and subtle components. These findings advance our understanding of the differences in the potential impacts of stroke on finger coordination, which may be useful in guiding targeted therapies to restore and enhance coordination complexity and expressiveness.

## Methods

### Participants

We analyzed data from 19 stroke survivors, 11 able-bodied older adults (OA), and 29 able-bodied younger adults (YA). Specifically, the stroke group comprised 12 survivors from a previously published dataset (<u>Xu et al., 2023</u>) and 7 newly enrolled chronic stroke survivors (30–80 years; 1 woman; Supplemental Table 1). The OA group comprised 11 newly enrolled able-bodied older adults (55–85 years; 5 women; 9 right-handed), and the YA (18-50 years) group comprised 29 able-bodied younger adults from the previously published dataset (<u>Xu et al.,</u> <u>2023</u>). Stroke diagnosis was confirmed by computed tomography, magnetic resonance imaging, or clinical report. Inclusion criteria were: 1) age ≥18 years, 2) ischemic stroke ≥6 months prior, 3) residual unilateral upper-extremity weakness, and 3) ability to provide informed consent. Exclusion criteria included 1) Fugl–Meyer Assessment >63/66, 2) MoCA <20/30, and 3) history of confounding neurological/orthopedic disorders, major psychiatric illness, or inability to perform the tasks. All participants provided written informed consent. Procedures were approved by the Johns Hopkins University and the University of Georgia Institutional Review Boards.

### Experimental Setup and Data Acquisition

#### Experimental setup and task

Experiments were performed and isometric forces were recorded using the Hand Articulation Neuro-rehabilitation Device (HAND; Carducci et al., 2002; Figure 1A-B). The HAND enables precise, independent measurement of isometric fingertip forces across all five fingers in three dimensions (3D). To use the HAND, participants were seated comfortably with the tested forearm wrist pronated and supported, and secured on the tabletop. Younger and older able-bodied participants (YA, OA) were tested using their dominant forearms, whereas stroke participants performed the task with their paretic and non-paretic forearms in separate blocks. Fingertips were secured in silicone cups custom-made to fit individual fingers and were mounted on adjustable force-sensing sticks. The forearm was supported by an ergonomic wristband and foams and secured with straps to minimize wrist and forearm movement. After the resting posture was adjusted to minimize baseline force (<1 N), the finger mounting sticks were locked in place, thereby fixing fingertip positions and preventing changes in hand posture across trials.

During individuated finger force production tasks, participants were instructed to produce isometric forces with one instructed finger to move a cursor in a 3D virtual space displayed on a computer monitor toward one of six virtual targets corresponding to three joint-space axes (MCP abduction/adduction, PIP flexion/extension, and MCP flexion/extension; Figure 1C). Fingertip forces were mapped to a Cartesian coordinate system in the virtual task space, in which the X-axis corresponded to MCP abduction (+X) and adduction (−X) within the frontal plane, the Y-axis corresponded to PIP flexion (−Y) and extension (+Y) within the sagittal plane, and the Z-axis corresponded to MCP flexion (−Z) and extension (+Z) within the sagittal plane. Participants controlled cursor movement along six directions (+X, −X, +Y, −Y, +Z, −Z) by producing isometric forces aligned with these joint motions.

Prior to this task, maximum voluntary force (MVF) for each finger-joint direction was determined using a range-of-motion task, in which participants generated isolated isometric forces while moving the cursor along each axis to their maximal voluntary force (with max force = 10 N). Each direction in the individuation task was then tested at four force levels corresponding to 20%, 40%, 60%, and 80% of MVF (see details in <u>Xu et al., 2023</u>). Each task was repeated four times. Participants were instructed to minimize non-instructed finger activation.

#### Data Acquisition and Preprocessing

Forces from all five fingertips were recorded simultaneously at 1000 Hz. Raw force recordings were low-pass filtered at 5 Hz (second-order Butterworth) and down-sampled to 10 Hz. For each trial, the force trace of the instructed finger along the target direction axis (e.g., index fingertip force along the X-axis; Figure 1D) was used to identify the primary force production cycle. Specifically, the time point of maximum force along the target axis was first identified. The trial segment was then defined as the interval between the nearest preceding and following local minima in that force trace, corresponding to the onset and termination of the force production cycle. This procedure ensured that exactly one force cycle was extracted from each trial, regardless of additional cycles in the force trace. In trials where the force produced by the instructed finger did not return to baseline after the peak, the end of the trial segment was defined using the minimum force velocity of the instructed finger immediately following the peak force. This extracted cycle captured the full rise and fall of the force trajectory for that trial. To enable reliable cross-participant and cross-task comparison, the trimmed data were then phase-aligned (see *Phase Estimation and Alignment* ) and baseline-centered (Figure 1D-F). To maintain consistent orientation across fingers, thumb forces were rotated 90° about the Y–Z plane, and left-hand data were mirror-flipped across the X-axis. All data analyses were done using custom-made MATLAB scripts (The MathWorks, Inc., 2024).

#### Phase Estimation and Alignment

To compare fingertip force data across trials, each trial was normalized based on progression through the movement cycle using phase estimation and alignment methods similar to those described previously (<u>Rosenberg et al., 2024</u>; <u>Rosenberg et al., 2026</u>). The instantaneous phase *ϕ*(*t*) was estimated as the four-quadrant phase angle from a phase portrait constructed using the recorded force signal in the instructed finger *x*(*t*).

To ensure a monotonically increasing phase estimate, the phase angle was unwrapped 2*π* and low-pass filtered using a fourth-order Butterworth filter (initial cutoff 6 Hz; iteratively reduced by 0.75 Hz until the phase became monotonically increasing, with a minimum cutoff of 0.25 Hz), and normalized from 0 to 2*π* for each trial (<u>Rosenberg et al., 2024</u>).

Each trial’s force data was then resampled onto a common set of evenly spaced phase values using a Von Mises kernel to perform phase alignment across trials (<u>Rosenberg et al.,</u> <u>2026</u>; <u>Winner et al., 2023</u>). The kernel weighted samples by their proximity to the target phase. The resampled force at each phase bin 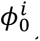 was computed as a weighted average:

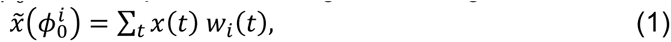

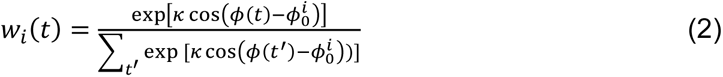

where 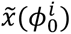 is the interpolated force value, *w*_*i*_(*t*) is the phase-dependent weight, and ***κ*** = 20 controls how strongly nearby phases are emphasized. This procedure yielded smooth, phase-aligned force trajectories that enabled comparison across participants regardless of force production speed or duration.

### Quantification of Flexor Bias Intrusion

Flexor bias was defined as the deviation of force trajectories toward flexion-dominated directions (−Y & −Z) (<u>Xu et al., 2023</u>). Intrusion of flexor bias due to stroke was defined as the difference in flexor bias between paretic and non-paretic fingers.

To assess whether increased flexor bias in the paretic hand relates to coordination patterns captured by high-variance components and to the loss of task and individual-specificity, we computed Pearson correlations to test whether intrusion of flexor-bias was associated with paretic vs. non-paretic differences in (i) reliance on high- vs. low-variance PCs, and (ii) coordination specificity, indexed by task- and individual-specific classification accuracy.

### Principal Component Analysis (PCA)

For each participant, 3D fingertip forces (F_x_, F_y_, F_z_) from all five fingers were concatenated across all tasks (5 instructed finger × 6 target direction × 4 force level). To examine spatiotemporal coordination complexity, PCA was applied individually to each participant across all tasks to extract spatiotemporal coordination patterns (<u>Yan et al., 2020</u>). Note that PCA was run in different manners for LDA classifications (see below).

#### Quantification of Coordination Complexity

1. To quantify finger force coordination complexity, we computed (1) the number of PCs needed to explain 95% of the variance in finger forces (variance accounted for; VAF) and (2) the variance explained by the first PC (VAF_1;_; <u>Schwartz et al., 2016</u>). A greater number of PCs required to explain 95% of the variance in each participant’s force trajectories suggests greater coordination complexity, whereas fewer PCs indicate lower complexity (<u>Santello et</u> <u>al., 1998</u>; <u>Tresch et al., 2006</u>; <u>Yan et al., 2020</u>).
2. The VAF_1_ metric, is a complementary continuous measure of complexity, in which lower VAF_1_ reflects a greater coordination complexity.

Both metrics were calculated independently for each participant, and group-level values were obtained by averaging across participants in each cohort: younger adults (YA), older adults (OA), stroke survivors’ non-paretic fingers (NP), and paretic fingers (Paretic). Group differences were evaluated using pairwise independent t-tests between groups.

#### Qualitative Assessment of High- and Low-Variance PC activations

To qualitatively assess differences between high- and low-variance PCs, we plotted individual-specific PC activations for PCs 1–3 (highest-variance) and PCs 13–15 (lowest-variance) derived from subject-level PCA of time-normalized isometric fingertip force trajectories (Fig. 3). Activations were visualized across representative tasks and instructed fingers to illustrate temporal coordination patterns captured by high- versus low-variance components.

#### Multidimensional Scaling and Isometric Force Trajectory Reconstruction from High- and Low- Variance PCs

To assess individual differences in the highest- and lowest-variance coordination components, we evaluated how well one participant’s PCs could reconstruct another participant’s fingertip force trajectories. Specifically, PC1 (highest variance) and PC15 (lowest variance) derived from each participant were used to reconstruct the force trajectories of all other participants. Reconstruction accuracy was quantified as the variance accounted for (VAF), producing a pairwise participant-by-participant VAF matrix. The resulting pairwise VAF matrix was treated as a similarity matrix and analyzed using multidimensional scaling (MDS; <u>Borg & Groenen, 2005</u>), such that greater distances in the MDS reflected greater differences in coordination between participants.

Isometric fingertip force trajectories were reconstructed using either the highest- (PC1) or lowest-variance (PC15) PCs. Reconstructions were obtained by multiplying the selected components by the corresponding temporal PC activations for each trial. Only PC1 or PC15 was used individually for reconstruction rather than the full set of PCs.

### Classification Analyses

#### Task-Specific Classification

To determine whether spatiotemporal fingertip force coordination patterns contained information about tasks, we performed linear discriminant analysis (LDA). PCA was performed in the same way as the spatiotemporal complexity analysis (see above). Task conditions were defined by the combination of instructed finger and target force direction. The PC activations were Z-scored and concatenated across time to form a feature vector for each trial with dimensionality (N_samples_ of PC activations)×(NPCs).

#### Individual-Specific Classification

Same as task specification, to distinguish among different individuals across tasks, time-varying PC activation trajectories were used. To ensure that individual differences were evaluated within a common coordination space, PCA was performed separately for each task using data across all participants.

For each trial, the resulting phase-varying PC activations were Z-scored and concatenated along the phase dimension to form feature vectors. These features were used as input to LDA classifiers, with participant IDs as the class labels. Classification was performed across tasks to assess whether spatiotemporal coordination patterns could reliably distinguish individuals independent of task condition within each group.

#### Group-Specific Classification

Same as task- and individual-specific classification, to distinguish among different groups (younger adult, older adult, non-paretic, paretic), phase-varying PC activations were used. PCA was performed separately for each task using data pooled across all participants. For each task, three-dimensional fingertip force trajectories from all five fingers were concatenated across participants and projected into a shared, task-specific PC space.

For each trial, the resulting temporal PC activation was Z-scored and concatenated along the phase dimension to form feature vectors. These features were used as input to LDA classifiers, with groups serving as class labels. This analysis assessed whether spatiotemporal coordination patterns contained information sufficient to distinguish impairment status across tasks.

To control for unequal group sizes, classification analyses for individuals and groups were performed on balanced datasets. Because the OA group contained the fewest participants (11 subjects), we used a bootstrapping procedure (1000 iterations) in which 11 samples from each impairment group were randomly selected, with replacement. The full classification pipeline, including PCA, feature construction, dimensionality reduction, and LDA classification, was executed independently for each iteration.

#### Evaluation of task-, individual- and group- specificity

To evaluate the ability of the remaining PCs to distinguish between different tasks, individuals, and groups, after iteratively removing higher-variance PCs in descending order of variance explained (Yan et al., 2020), LDA classifiers were trained using PC activations after those of the first N PCs (i.e., the N components explaining the most variance in isometric finger force trajectories) were excluded from the feature set. For example, when N = 0, the activation profiles of all PCs were included in the feature set. When N = 1, the activation profile for PC1 was excluded from the feature set, and the classifier was trained and tested using only the activations of PC2 through PC15. This procedure was repeated until only the lowest-variance PCs activation patterns (i.e., PC15) remained in the feature set.

Classification accuracy was assessed using leave-one-task-out cross-validation: For each task condition (defined by instructed finger and force direction), one trial among four repetitions was held out as the test sample while the classifier was trained on the remaining three trials. This procedure was repeated so that each of the four trials served once as the test set. Classification accuracy was averaged across folds and 1000 bootstrap iterations.

#### Null Models for Chance-Level Performance

To establish chance-level baselines for LDA classification analyses, we trained null models by repeating all LDA classification analyses with permuted class labels. For each analysis (task-, individual-, and group-specificity), class labels were randomly shuffled across trials while preserving the same number of trials per class. LDA models were then trained and tested using the same cross-validation procedure as the true data. This permutation procedure was repeated 1,000 times to generate a null distribution of classification accuracies for each analysis. Observed classification accuracies from the real data were then compared against the corresponding null distribution using independent-samples t-test to determine whether performance exceeded chance level. Statistical significance was assessed by computing empirical p-values as the proportion of null accuracies greater than or equal to the observed accuracy.

### Statistical analyses

#### Linear Mixed Effects Models

Linear mixed-effects (LME) models were used to assess how coordination complexity and classification-based specificity varied as a function of PC Index (treated as a categorical factor), Group (YA, OA, NP, Paretic), and Measurement Type (endpoint vs. spatiotemporal).

To assess spatiotemporal coordination complexity, an LME model was fit to VAF values derived from participant-level PCA. VAF values for all 15 PCs were analyzed with fixed effects of Group (YA, OA, NP, Paretic), PC Index (treated as a categorical factor), and their interaction, and a random intercept for Participant to account for repeated measurements across PCs within participants. The YA group and PC1 served as reference levels.

To determine whether task-, individual-, and group-specific classification accuracy varied as a function of group and the progressive removal of higher-variance PCs, LME models were fit to classification accuracy values computed for each step of PC removal. Fixed effects included Group, PC Index (treated as a categorical factor), and their interaction. Random intercepts were included for Participant when classification accuracy varied across individuals and for Task when accuracy varied across tasks, accounting for repeated measurements across each PC Index. The YA group and the condition with all PCs included served as reference levels. A separate model was fit for each classification (task, individual, and group-specificity).

To compare endpoint and spatiotemporal force representations, LME models were fit to classification accuracy values with fixed effects of Measurement Type, PC Index, and their interaction. A random intercept for Observation was included to account for repeated measurements across PC indices within the same replicate. Endpoint measures and the condition with all PCs included served as reference levels. A separate model was fit for each group and each classification.

Statistical significance of fixed effects and interactions was assessed using Type III analysis of variance with Satterthwaite approximation for degrees of freedom.

#### Independent samples t-tests

To determine whether classification performance exceeded chance, we compared observed classification accuracies with permutation-derived null accuracies generated by randomly shuffling class labels using independent-samples t-tests. Independent-samples t-tests were also used to assess group differences in angular deviation, number of PCs required to reach 95% VAF, and classification accuracy when all PCs were included. Independent samples t-tests were used to evaluate differences in task-, individual-, and group-level classification accuracy between force magnitudes (20% vs 80% MVF).

#### Pearson Correlations

To assess associations between flexor bias intrusion and classification accuracy for tasks, individuals, and groups, we conducted Pearson correlation analyses. Pearson correlations were also used to evaluate associations between flexor bias intrusion in paretic fingers and the variance in isometric fingertip force trajectories captured by PC1 and PC15.

#### Chi-square tests of independence

To evaluate whether higher- and lower-variance PCs differed in their contribution to classification performance, chi-square tests of independence were used to compare the frequency with which higher-variance (PC1-7) and lower-variance (PC8-15) components contributed significantly to group classification.

### LDA Coefficient Analysis

To assess the statistical reliability of the LDA coefficients, we used a nonparametric bootstrapping procedure. For each pairwise LDA classification, trials were resampled with replacement, and the LDA model was refit on each resampled dataset (10,000 bootstrap iterations). This procedure generated a bootstrap distribution of coefficient values for each PC. For each PC, a 99% confidence interval (CI) was constructed from the bootstrap distribution. PCs whose confidence intervals did not include zero were considered to contribute reliably to classification performance. To visualize how they contributed to finger forces, each PC identified as significant during bootstrapping was separately used to reconstruct three-dimensional fingertip force trajectories using its coefficients and activations.

To evaluate whether dominant PCs contributed more often to classification than lower-variance PCs, the entire set of PCs was divided into higher-variance (PC1–7) and lower-variance (PC8–15) groups. We then tallied the number of PCs that significantly contributed to each classification type (task, individual, impairment). Differences in contribution frequency between higher- and lower-variance PCs were evaluated using Chi-square tests of independence. All statistical analyses us a significance level of α = 0.05.

### Synthetic Data Generation and Projection

To assess an alternative explanation that the above-chance accuracy of classification using only lower-variance PCs (e.g., PC15) may be due to artifacts such as sensor noise, we generated synthetic 14-dimensional signals across 120 classes (tasks), matching the number of task conditions in the in vivo dataset. Each class contained 100 samples, approximating the number of samples per task in the experimental isometric fingertip force recordings. Samples were constructed as random mixtures of sinusoidal or cosinusoidal basis functions modulated by smooth envelopes. Random linear weights, amplitude scaling, and baseline shifts were applied to produce distinct yet systematically related class signals. To mimic repeated conditions in the experiment, the dataset was duplicated across four repetitions.

The complete dataset was then embedded in a 15-dimensional space via multiplying the 14-dimensional data matrix by a random Gaussian mixing matrix:

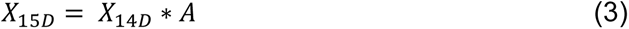

where X_14*D*_ ∈ ℝ^*Nx*14^ is the concatenated dataset, and A ∈ ℝ^14*x*15^ is a Gaussian random mixing matrix.

Gaussian noise was added to the projected data at four levels (σ = 0%, 1.0%, 2.0%, and 5.0% of the maximum signal amplitude). For each noise condition, PCA was applied to extract 15 PCs, and cumulative VAF was computed. The same PC-removal and LDA classification procedure used to assess the task-specificity of the experimental data was then applied to the synthetic dataset. Specifically, higher-variance PC activations were iteratively removed and the remaining lower-variance PC activations were used as input features for LDA classifiers to predict task identity. Classification accuracy obtained from the lowest-variance components (e.g., PC15) was evaluated to determine whether additive noise alone could produce above-chance classification performance.

## Supplemental Figures

**Supplemental Table 1.** Patient characteristics. Data indicate patients’ gender (M, male; F, female), age (years), time since stroke (in months), handedness (L, left; R, right), affected side (L, left; R, right), Montreal Cognitive Assessment (MoCA, maximum 30), Fugl-Meyer Assessment for upper extremity (FMA, maximum 66), and Action Research Arm Test (ARAT), and grip strength (in pounds) if applicable.

| Patient | Gender | Age | Months post stroke | Handedness | Affected side | MoCA | FMA | ARAT | Grip strength (affected) | Grip strength (unaffected) |
| --- | --- | --- | --- | --- | --- | --- | --- | --- | --- | --- |
| SP001 | F | 30 | 60 | R | L | 27 | 51 | 37 | 39 | 72 |
| SP002 | M | 67 | 112 | R | L | 28 | 20 | 6 | 28 | 106 |
| SP003 | M | 70 | 61 | R | L | 27 | 20 | 17 | 17 | 83 |
| SP004 | F | 45 | 16 | R | L | 24 | 40 | 39.5 | 10.5 | 43.5 |
| SP005 | M | 51 | 83 | R | L | 20 | 43 | 40 | 31 | 133 |
| SP006 | M | 59 | 63 | R | L | 24 | 34 | 9 | 18 | 93 |
| SP007 | F | 64 | 119 | R | L | 26 | 23 | 4 | 27 | 63 |
| SP008 | F | 69 | 97 | L | R | 29 | 26 | 16 | 25 | 75 |
| SP009 | M | 80 | 36 | R | L | 28 | 58 | 56 | 45 | 53 |
| SP010 | F | 68 | 33 | R | R | 26 | 56 | 57 | 43 | 45 |
| SP011 | F | 64 | 3 | R | R | 25 | 58 | 57 | 50 | 67 |
| SP012 | M | 49 | 95 | R | L | 30 | 19 | 7 | 44 | 157 |
| SP013 | F | 69 | 97 | R | R | 27 | 64 | 57 | 58 | 65 |
| SP014 | M | 78 | 8 | R | R | 26 | 56 | 46 | 22.7 | 49.3 |
| SP015 | M | 68 | 27 | R | L | 27 | 39 | 40 | 28.3 | 92.3 |
| SP016 | M | 63 | 9 | R | R | 27 | 62 | 57 | 79.3 | 83.7 |
| SP017 | F | 51 | 14 | R | L | 30 | 62 | 57 | 37 | 42 |
| SP018 | M | 56 | 43 | R | R | 29 | 40 | 47 | 17.7 | 32.3 |
| SP019 | M | 63 | 3 | R | L | 26 | 44 | 41 | 24 | 56 |
| SP020 | M | 81 | 28 | R | R | 24 | 61 | 45 | 41.7 | 50.7 |

**Supplemental Figure 1:**
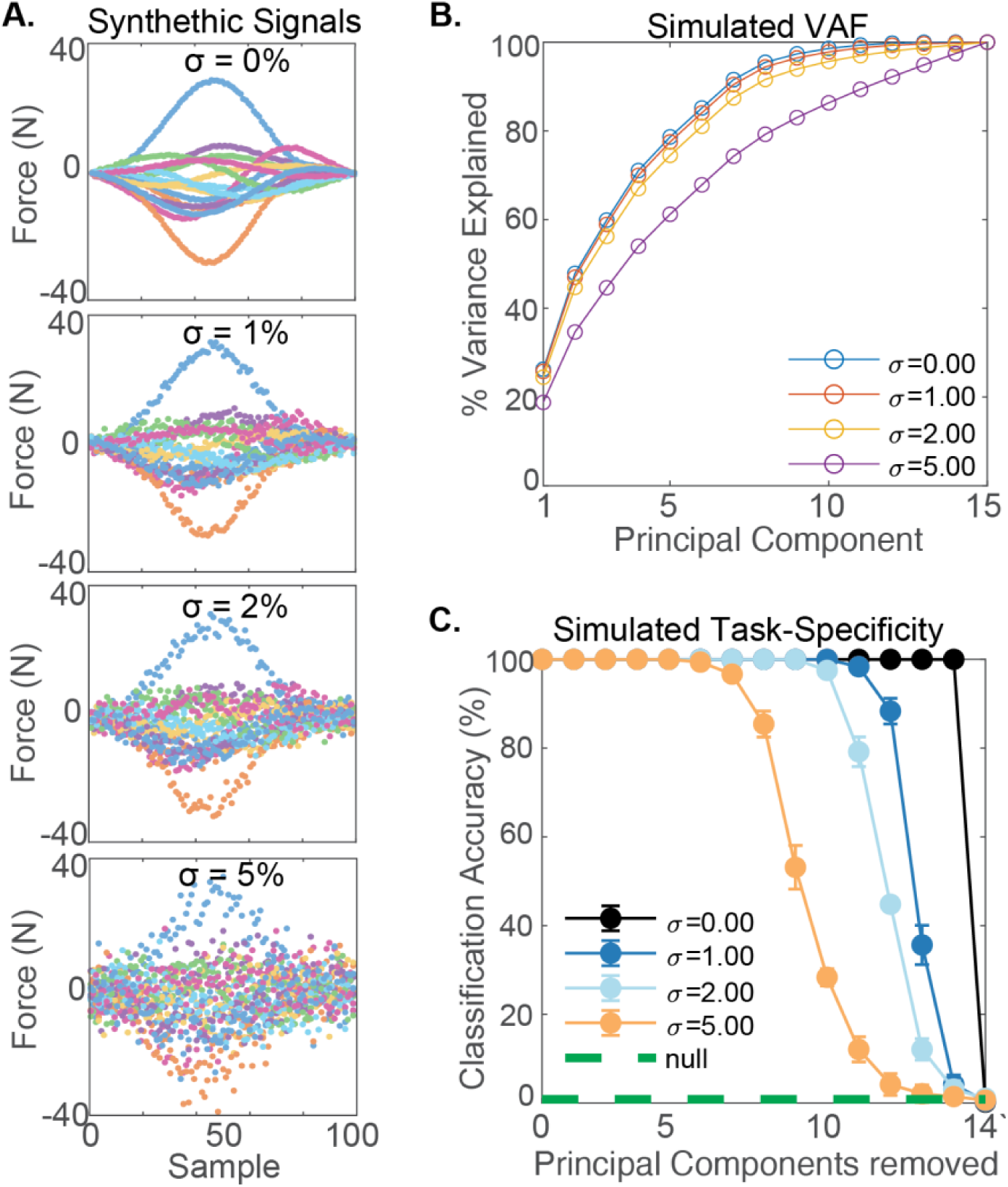
Simulated signals, variance explained, and task-specificity under increasing noise conditions. **(A)** Example simulated 14-dimensional signals at four noise levels (σ = 0, 1, 2, 5). At σ = 0, signals show smooth oscillatory temporal patterns, while higher noise progressively distorts temporal patterns. Signals were projected onto a 15^th^-dimension using random matrix mixture. **(B)** % The cumulative VAF explained by the first 15 principal components (PCs) is averaged across simulations and shown across noise levels. In the noise-free condition, more than 95% of the variance in the synthetic signals is accounted for by the first approximately 10 PCs. **(C)** Task-specific classification accuracy after gradual removal of the high-variance PCs. Accuracy remained near 100% in the noise-free condition until nearly all PCs were removed, whereas classification performance degraded more rapidly under increasing noise. The dashed green line suggests chance-level performance.

**Supplemental Figure 2.**
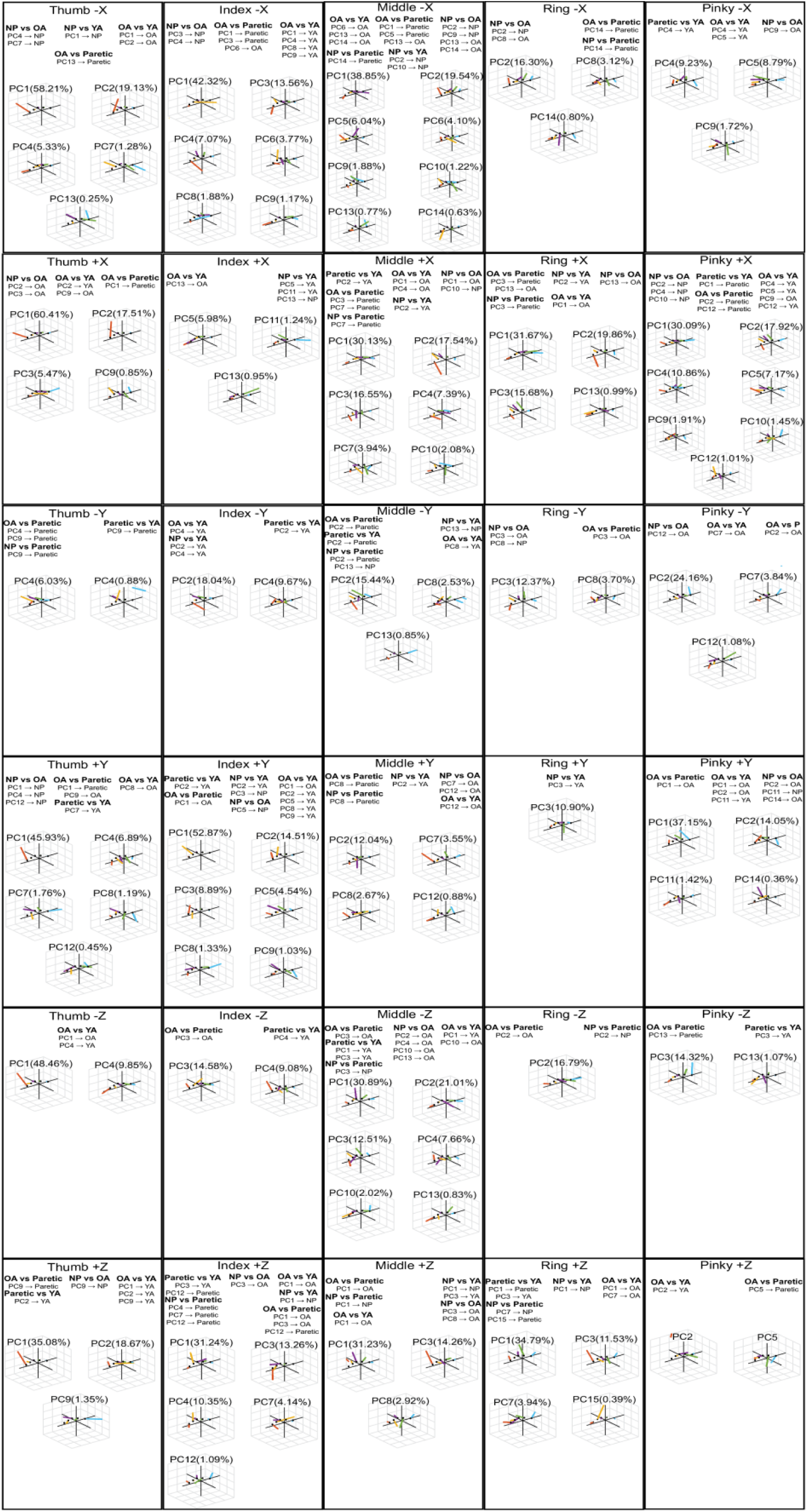
Three-dimensional representations of significant principal components contributing to pairwise group classification. Each cell represents a specific task, with rows organized by instructed finger and columns organized by force target direction (+/–X, Y, Z). Within each cell, the 3D representations of PC coefficients show spatiotemporal coordination patterns which significantly contribute to distinguishing pairs of groups (younger adults, older adults, non-paretic, and paretic hands). Significant PCs were identified using bootstrap-derived 99% confidence intervals on LDA coefficients (see Methods).

**A library of spatiotemporal finger coordination for characterizing able-bodied and post-stroke finger coordination.**

Current assessments of post-stroke finger impairment often emphasize coarse measures such as strength, grasp success, range of motion, or spasticity (<u>Fugl-Meyer et al., 1975</u>; <u>Lyle, 1981</u>; <u>Wolf et al., 2001</u>). In contrast, our approach allows us to identify the coordination patterns that drive pairwise group classification across different tasks, allowing us to identify a catalog of spatiotemporal components that characterize YA, OA, NP, and paretic finger coordination. Using this catalog, we capture detailed spatiotemporal patterns that distinguish both able-bodied and impaired coordination. Summarized in Supplemental Figure 2, this catalog provides a richer description of stroke-impaired finger coordination than those typically available in clinical assessments and highlights the potential of this framework for personalized assessment and rehabilitation, enabling comparison of an individual’s coordination patterns against group-specific coordination patterns to assess whether their coordination patterns are similar to those of paretic fingers.

