## Supplementary figures and images for "Loss of hand control expressiveness revealed by task- and individual-specificity in spatiotemporal finger coordination"

### Supplemental Figure 1

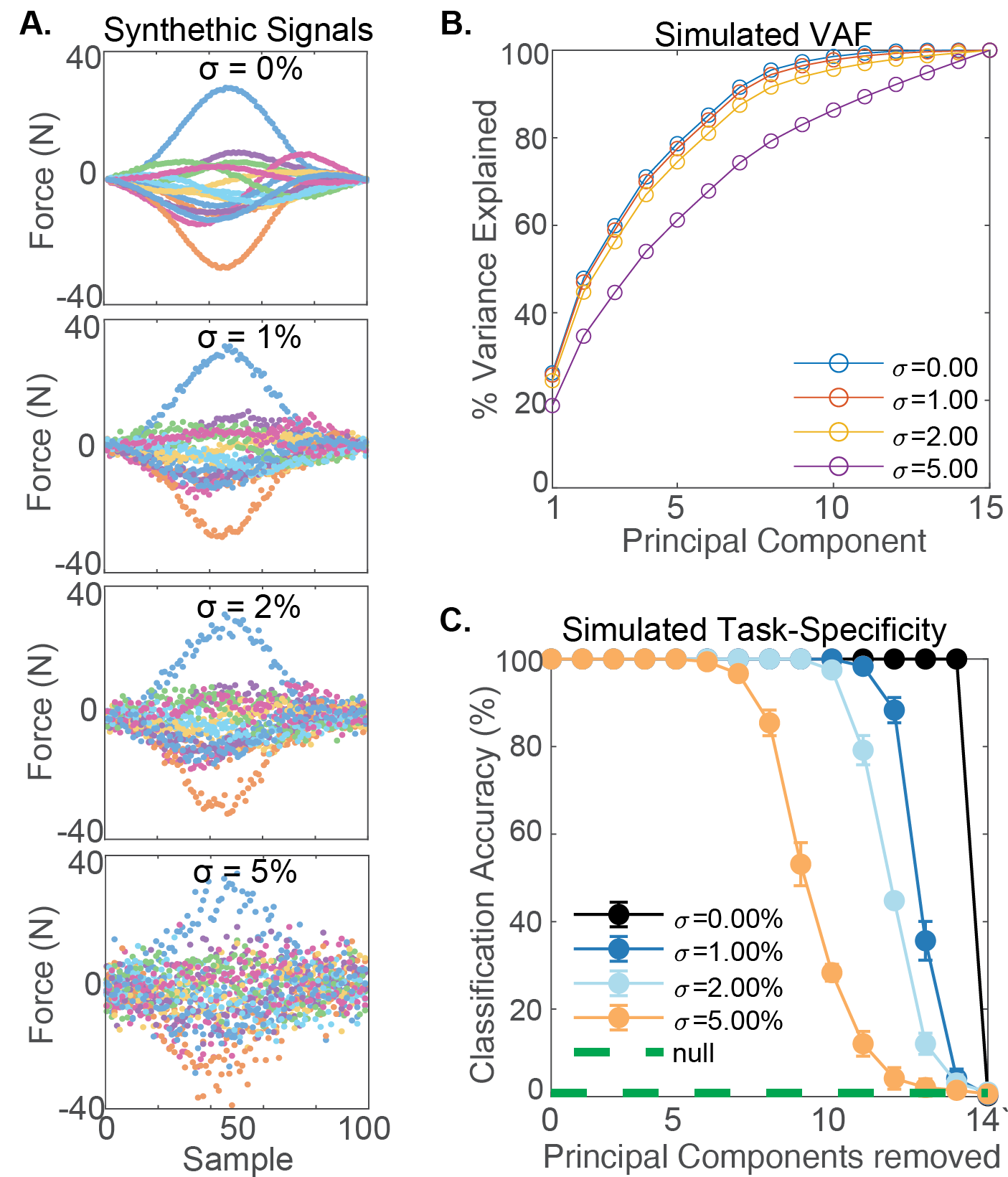

### Supplemental Figure 2

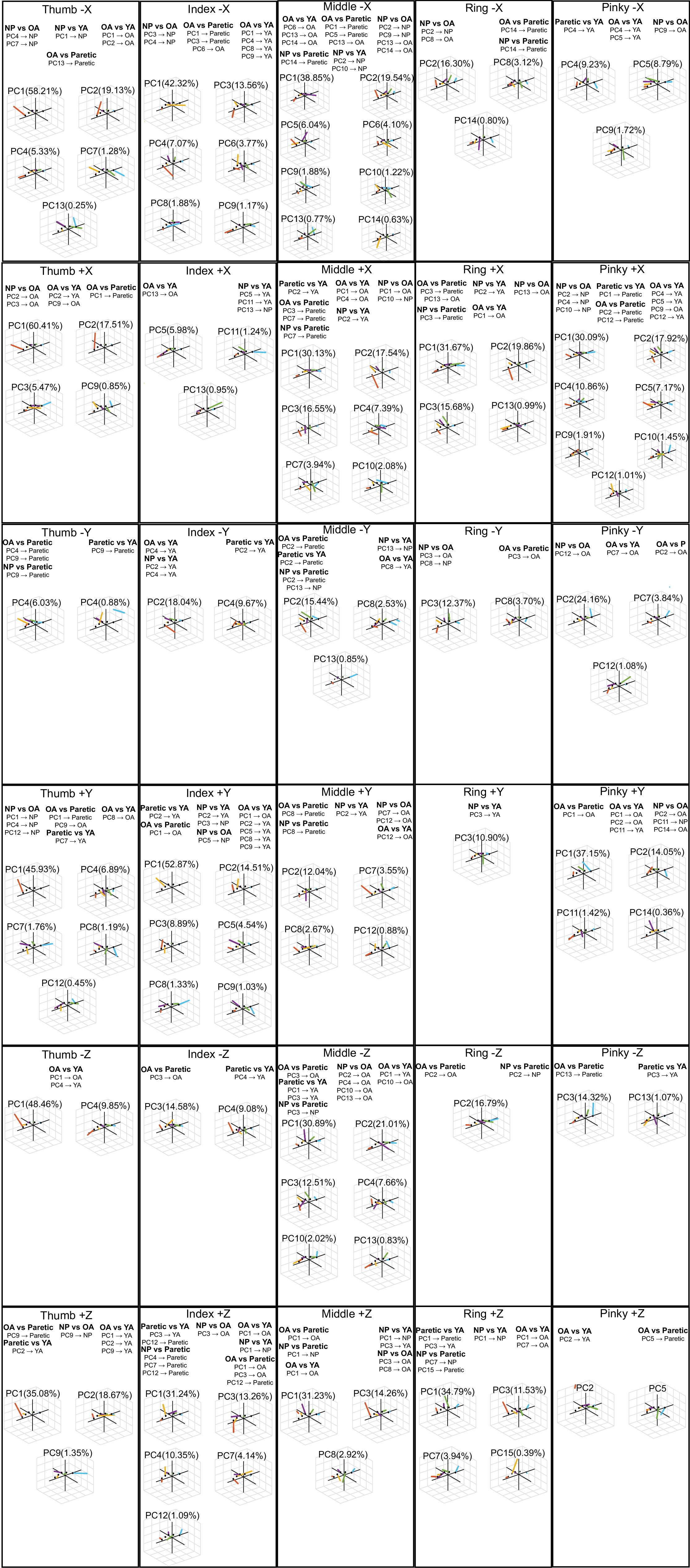

### Supplemental Table 1

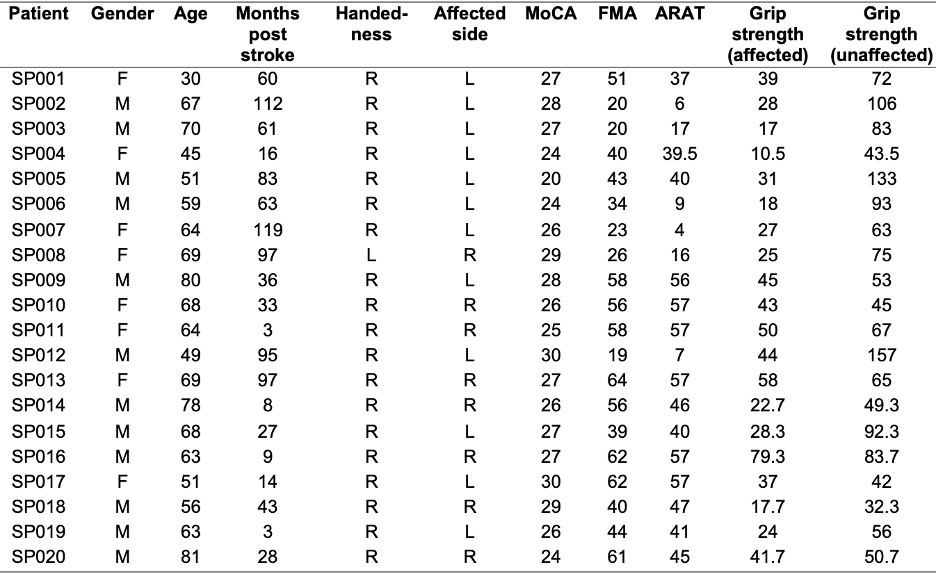
